# Estimation of adult census size from close-kin dyads in the malaria mosquito *Anopheles gambiae*

**DOI:** 10.64898/2026.04.01.712345

**Authors:** Anita Lerch, Scott T. Small, Tin-Yu J. Hui, Priscila Bascuñán, Ellen M. Dotson, Martin Lukindu, Krystal Birungi, Jonathan K. Kayondo, Austin Burt, T. Alex Perkins, Nora J. Besansky

## Abstract

Accurate estimates of adult mosquito abundance are central to the design and evaluation of vector control strategies, yet they are difficult to obtain from natural populations. Conventional mark-recapture methods for estimating adult mosquito census size pose logistical and other challenges. A recently developed close-kin mark-recapture (CKMR) approach is a promising alternative. However, application of CKMR has been largely confined to long-lived vertebrates. Validation on empirical data from short-lived, highly fecund insects is lacking. Here, we apply CKMR to a natural population of *Anopheles gambiae*, the primary African malaria mosquito, sampled from a small island in Lake Victoria, Uganda. Using a high-diversity amplicon panel of genome-wide markers, we genotyped 714 adult mosquitoes collected over a 20-day period. We classified pairs to close-kin categories by implementing probabilistic latent-kinship estimation rather than using a deterministic threshold-based framework. We observe numerous full-sibling pairs but no parent-offspring pairs, a pattern indicative of extreme variance in reproductive success owing to frequent failure of mosquito egg clutches to produce adults. To adapt the CKMR framework to mosquito life history, we explicitly modeled the mosquito life cycle including the possibility of clutch failure. By incorporating probabilities of both parent-offspring and full-sibling relationships, we estimate an adult female census size of 26,887 (95% credible interval: 6,979 - 146,011), and a clutch failure probability of 97.6% (CrI: 91.3 - 99.6%). We use these estimates to calculate the predicted variance in reproductive success and effective population size. Through individual-based simulations we confirm our estimates and the necessity of modeling reproductive variance to explain observed kinship patterns. Our results demonstrate that CKMR can be applied successfully to mosquitoes, provided that appropriate adjustments are made to account for their natural history.

## 1. Introduction

Census size—the number of adult individuals in a population—is a fundamental parameter in basic and applied ecology (Krebs 2009). Estimates of population abundance are important for understanding population and ecosystem dynamics. In applied contexts, they inform management of threatened or threatening species and help to guide interventions by providing a baseline for evaluating population-level change. Despite its importance, census size remains difficult to estimate for many organisms, particularly those that are super-abundant, highly mobile, or difficult to detect in the landscape and oceans.

Nowhere is the need more evident than for arthropod disease vectors. Planning and evaluation of vector control interventions requires feasible and reliable methods to estimate and track abundance. In particular, the successful deployment of gene drive systems and other genetic control strategies that aim to suppress vector populations and reduce disease transmission will depend on accurate estimates of population size to determine release ratios and interpret post-release dynamics (Dhole et al. 2020; Rasic et al. 2021; Hancock et al. 2024). In Africa, where malaria remains a major public health burden, the need is particularly acute for *Anopheles gambiae sensu lato*, the primary vectors responsible for most malaria transmission (World Health Organization 2024).

Historically, mark-release-recapture (MRR) has been the dominant method for estimating animal abundance (Seber 1982). Although MRR has been applied widely to mosquito populations (Guerra et al. 2014), it presents substantial logistical challenges. MRR studies typically require the release of large numbers of marked laboratory-reared mosquitoes, which may differ behaviorally and genetically from wild individuals. Recapture rates are often low, reducing statistical precision.

Close-kin mark–recapture (CKMR) is a recent and powerful methodological advancement that avoids the pitfalls of MRR (Bravington, Skaug, and Anderson 2016). Rather than physically marking and recapturing individuals, CKMR uses multilocus genotyping to identify close relatives within a population sample. Pairs of close kin act as the genetic analogs of capture-recapture events, with the expected frequency of such pairs declining as population size increases (Bravington et al. 2016). Initial CKMR methodology was tailored for aquatic vertebrates, particularly commercially exploited fish, that were its exclusive subjects until recently. As interest in the method has expanded, especially in the conservation and wildlife management realm, the CKMR framework has been extended to terrestrial vertebrate systems (Lloyd-Jones et al. 2023; Merriell et al. 2024). However, the robustness of its performance for demographic analysis of short-lived, highly fecund insects remains untested.

CKMR has potentially broad taxonomic scope for sexually reproducing species, with few exceptions (Bravington et al. 2016). Wider implementation depends on accommodating the biology of the target species in both estimation models and sampling design, taking into account complexities such as variation in expected kin relationships and reproductive output (Merriell et al. 2024). Fundamental to CKMR is accurate categorization of close-kin relationships, determined from genetic data. Yet, certain kinship relationships (half-siblings, grandparent-grandchild, and avuncular) are impossible to distinguish genetically with unlinked markers alone, as they have the same expected number of gene copies that are identical-by-descent (Bravington et al. 2016)(Table S1). For long-lived and late-maturing vertebrate species whose mating structure approximates random, the expected frequency of confounding kin types is often low but can be further minimized by judicious sampling design (*e*.*g*., controlling for age, life stage, geographic location) or by eliminating modeling of the confounded kin relationships altogether and including only unambiguous close-kin categories (*e*.*g*., parent-offspring). For highly fecund mosquitoes, where adult lifespan is short, generations overlap, and females mate once but are iteroparous, the expected frequency of temporal and spatial overlap of difficult-to-distinguish kinship categories is substantial. To date, the only extension of the CKMR framework to mosquitoes using empirical data has been the development of spatially explicit models aimed at estimating dispersal in *Aedes* (Jasper et al. 2019; Filipovic et al. 2020; Schmidt et al. 2021; Schmidt et al. 2023). In most of these studies, the problem of confounding kin types was controlled by sampling only immatures from deployed ovitraps over the span of a single generation, conditions expected to yield only full-sibling, half-sibling, and cousin kinships (Schmidt et al. 2023). Recent work has established a theoretical framework for adapting CKMR to *Anopheles* mosquito demography (Sharma et al. 2022), but because this was an *in silico* study, it assumed perfect kinship inference.

Field studies of *An. gambiae* ecology show that juvenile recruitment is concentrated in a small and temporally variable subset of aquatic habitats, with most oviposition events failing due to abiotic or biotic conditions (Mutuku et al. 2006; Munga et al. 2007). Although recognized by field biologists, this phenomenon has rarely been quantified and has underappreciated consequences for *An. gambiae* population demography. Such dynamics imply substantial variance in realized reproductive success among adults. If high reproductive variance is not accounted for in the CKMR estimation model, abundance estimates could be biased. Here, we apply CKMR to a natural population of *An. gambiae sensu stricto* (hereafter, *An. gambiae)* sampled from Jaana, a small island in Lake Victoria, Uganda. Our goal was to determine whether CKMR can yield informative estimates of mosquito adult abundance and, if so, what modifications are required to make such inference biologically realistic. We developed a high-diversity genotyping panel of genome-wide markers and used it to infer close-kin relationships among 714 field-collected adults. We accounted for uncertainty in kinship assignments by treating true relationships as latent variables within a pseudo-likelihood CKMR framework, explicitly propagating uncertainty in relationship inference.

We detected numerous full-sibling pairs but no parent-offspring pairs. We show that this pattern is explained by frequent failure of mosquito egg clutches to produce adults. By incorporating a clutch failure parameter into the demographic model, we obtain stable estimates of adult female census size that are consistent with both observed kinship patterns and individual-based simulations. We also estimate variances of female and male reproductive success and the effective population size. Together, these findings establish CKMR as a robust and biologically realistic tool for estimating mosquito abundance with direct relevance to ecological studies and applied vector control strategies.

### 2. Materials and Methods

### 2.1 Amplicon Genotyping Panel and Evaluation using Archived Island Samples

We used the Uganda-specific variant call format (VCF) file from Ag1000G phase I (2017) to identify SNPs segregating in the population at ≥5% (Miles et al. 2017). Candidate microhaplotype loci containing ≥3 variable SNPs in a 100-bp span were chosen to cover all chromosome arms, excluding known repetitive, heterochromatic, and inversion regions (2La, 2Rb) in the *An. gambiae* genome (coordinates provided in Table S11 of Bergey et al. (2020)). Primers flanking candidate loci were designed using a multiplex primer design pipeline (Campbell et al. 2015), resulting in 355 loci passing all computational filtering steps in the pipeline.

DNA was extracted using a CTAB method (Chen et al. 2010) from archived *An. gambiae* specimens previously collected in 2015 from two Ssese Islands in Lake Victoria (Lukindu et al. 2018; Bergey et al. 2020): Bukasa, N=51; Sserinya, N=43. We genotyped these specimens using Genotyping-in-Thousands by Sequencing (GT-Seq) (Campbell et al. 2015). Briefly, individually indexed multiplexed PCR products were normalized for DNA concentration, pooled, and sequenced on a MiSeq instrument using 2 × 75 paired-end cycles. Initial genotyping by GTseek LLC with 355 *An. gambiae* markers followed the GT-seq protocol (Campbell et al. 2015). The initial test sequencing run revealed primers that produced an overabundance of artifact sequences during multiplex PCR. After removing these primers, a second sequencing library was constructed and sequenced, resulting in a final set of 354 compliant loci (Appendix 1).

Sequencing reads were grouped by index and processed bioinformatically. After adapter removal, reads were mapped to the *An. gambiae* genome (AgamP4) and retained only if they mapped uniquely to the expected chromosome. Next, the paired-end reads from each locus were merged using NGmerge (Gaspar 2018) with minimum and maximum overlaps of 10-bp and 100-bp. Successfully merged reads were retained and aligned using BWA-MEM (Li and Durbin 2009) to a reference file of consensus amplicon sequences derived from the AgamP4 reference genome. Using BCFtools (Danecek et al. 2021), SNPs at each locus were called using a pooled approach in which all individuals are analyzed jointly as one population. The VCF file output by BCFtools was filtered to retain only biallelic SNPs with a minimum depth of 20, and a minimum quality of 30, at a locus within an individual. Sites missing data from more than 20% of individuals in the sample were removed, as were indels.

We used MICROHAPLOT (Ng et al., https://doi.org/10.5281/zenodo.820110) to assemble phased microhaplotypes from each individual from variants on single reads, and to perform further filtering following Baetscher et al. (2018). We removed haplotypes with less than 20 reads at a locus within an individual. Loci for which this minimum read depth was violated in more than 75% of individuals were removed. In addition, loci for which a minimum read depth of 50 resulted in more than two haplotypes within an individual, owing to sequencing errors and ‘index switching’ (Sinha et al. 2017), were removed. For those remaining, genotypes were called. Homozygotes were defined as individuals with one haplotype. An individual was defined as a heterozygote only if the two haplotypes were in the bounds of expected 1:1 allelic balance, defined as a ratio of sequencing reads for the first and second alleles >0.6. Individuals deviating from this ratio were defined as homozygotes. If more than 15% of potentially heterozygous individuals in the sample deviated from the expected allelic balance at a locus, that locus was excluded from the panel. Finally, loci that deviated from Hardy-Weinberg equilibrium, as well as those that were monomorphic (homozygous for the same haplotype across the entire population sample), were removed. The remaining loci constitute the 291-locus genotyping panel used in this study (Appendix 2).

### 2.2 Controlled Crosses of *An. gambiae* G3 to Generate Known Kinships

The G3 colony is maintained in the insectary at the Centers for Disease Control and Prevention as part of the BEI Resources Malaria Research and Reference Reagent Research Center (MR4). We designed a sequential series of single-pair matings, conducted as described (www.beiresources.org/Publications/MethodsinAnophelesResearch.aspx). The crossing scheme (Fig S1) created a pedigree with known kinship relationships, including parent-offspring, full-sibling, half-sibling, full-cousin, and half-cousin.

DNA was extracted from the carcasses of all 15 parents in generations 1 and 2, and 15-20 male and female full-siblings from eight different families at generation 3. In addition, DNA was extracted from 20 ‘unrelated’ mosquitoes from this colony (Fig S1). As no pair of mosquitoes from an inbred colony under long-term laboratory culture is completely unrelated (Norris et al. 2001; Lainhart et al. 2015), this design provides a conservative test of power for kinship inference. GT-Seq was performed with the 291-locus genotyping panel to determine individual genotypes.

### 2.3 Inference of Simulated and Known Kin-Pairs Using the Standard Classification Threshold Method

We simulated genotype pairs of ‘known’ kin categories both for the island samples and the G3 colony as follows. Using the R program CKMRsim (Anderson 2020), we estimated allele frequencies from genotype frequencies. For the G3 pedigree, this estimation was performed using only the unrelated specimens: the set of 20 unrelated G3 mosquitoes, together with all maternal parents and the single father in generation 1 (Fig S1). This ensured that any allele carried into generation 3 sibships will have been accounted for by CKMRsim. With the ‘simulate_Qij’ function of CKMRsim, allele frequencies at each locus were used as the basis for simulations of 100,000 multilocus genotype pairs of individuals for each of the ‘known’ relationships of interest (parent-offspring, full-sibling, half-sibling, and unrelated). We assumed that markers on the same chromosome are linked [unlinked=F, with distance between marker adjusted with the average recombination rate of 1.27 per Miles et al. (2017)], and we accounted for genotyping error (per-locus, miscall_rate=0.005, dropout_rate=0.005, per_snp_rate=F).

From the simulated genotypes, per-pair log-likelihood ratios were calculated of one relationship versus another. The distributions of simulated log-likelihood ratios were examined to identify a log-likelihood ratio threshold for classifying a pair into a given relationship (parent-offspring, full-sibling, half-sibling, unrelated). This threshold determines the false negative rate (FNR; the per-pair rate at which a pair truly sharing a particular relationship is misclassified as something else) and the false positive rate (FPR; the per-pair rate at which a pair is falsely classified into a particular relationship). Because the FNR and FPR have an inverse relationship, a choice must be made about how to balance these distinct types of classification error when specifying the threshold. To address this, we chose to maximize the overall classification accuracy (ACC), which is defined as

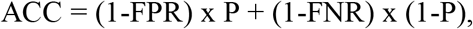

where P is the proportion that truly have a given kinship type. In simple terms, the overall classification accuracy is the proportion of pairs that are correctly classified as being in one kinship category as opposed to another. The proportion P is expected to differ based on the kinship category, population size, and other factors. To ensure that we worked with biologically plausible values of P for each kinship category, we computed values of P for each kinship category using outputs of the simulation model (see Methods 2.7).

In the case of the G3 pedigree, we used a random subset of known parent-offspring, full-sibling, and half-sibling pairs from the genotyped pedigree to assess whether the known pairwise relationships would be accurately classified by this method.

### 2.4 CKMR Sampling on Jaana Island

Jaana is an island 3-4 km long and 1-2 km wide, located in the Ssese Island archipelago of Lake Victoria in the Kalangala District of southeastern Uganda (Fig 1). Jaana hosts three small villages of ~60 to ~370 houses, from which indoor resting adult mosquitoes were collected daily during 20 days, from 23-October to 11-November-2021, except for three days of adverse weather or community events. Adult mosquitoes were collected indoors between 6-8 AM with Prokopak aspirators (Vazquez-Prokopec et al. 2009), from consenting households. To achieve balance and avoid excessive repeat sampling of the same households, LWZ and KKU were divided into adjacent sections: LWZ-A (~200 houses), LWZ-B (~70 houses), LWZ-C (~100 houses), KKU-A (~60 houses), and KKU-B (~50 houses). NKD, consisting of only ~60 houses, was not subdivided. Randomly selected consenting households from the relevant villages and sections were sampled in a repeating 3-day rotation: LWZ-A (day 1), LWZ-B+C (day 2), KKU-A+KKU-B+NKD (day 3). The mean number of adult *An. gambiae* collected daily was 43, and the total number collected over the 20 days was 733, the overwhelming majority from LWZ (Table S2).

**Figure 1.**
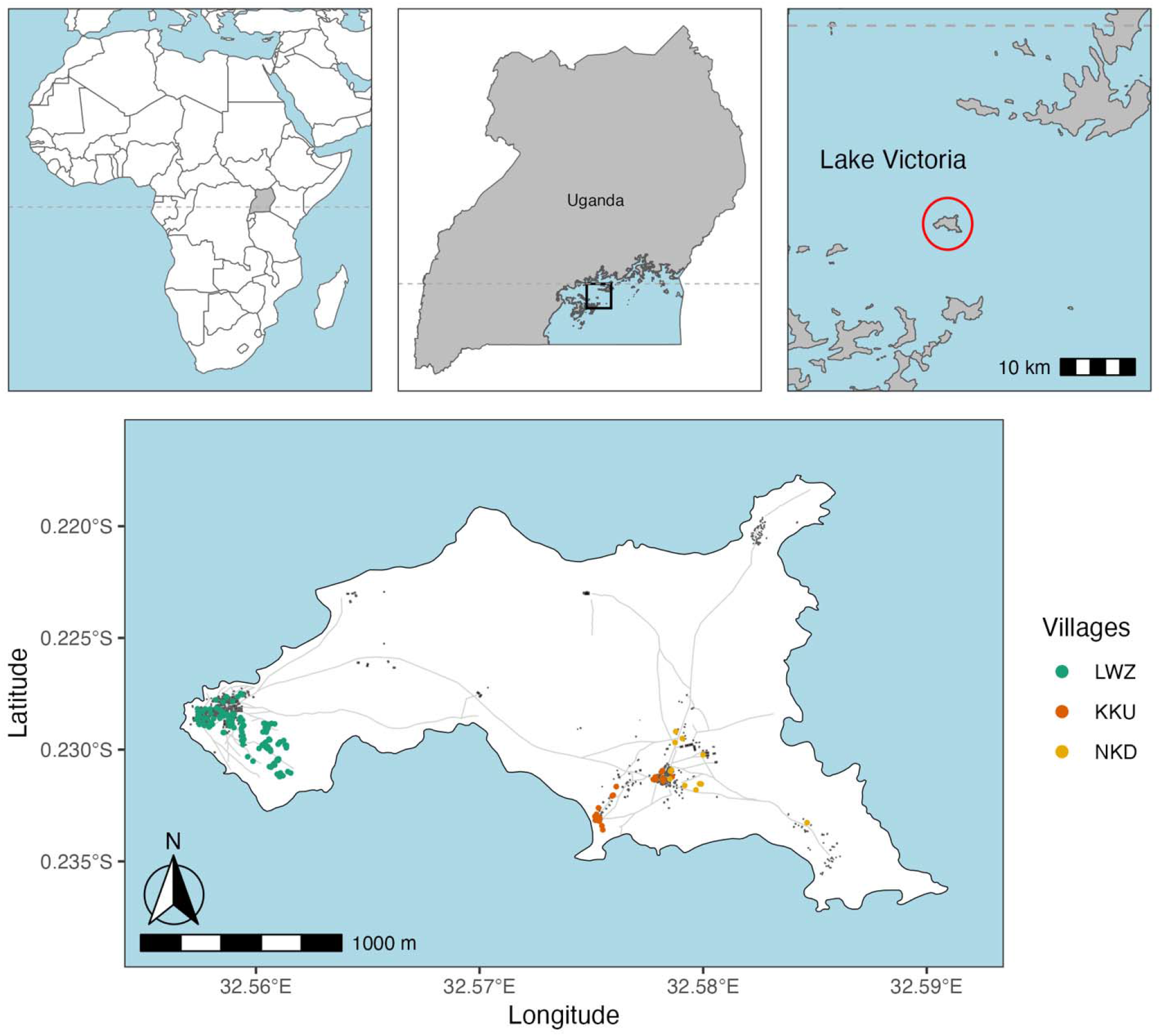
Geographic location of Jaana Island and its sampling locations. Top (left to right): Uganda within Africa (gray shading); Ssese archipelago in northwestern Lake Victoria offshore Uganda (black box); Jaana Island within the Ssese archipelago (red circle). Bottom: Households with mosquitoes collected in the villages of Lwazzi (LWZ), Kikku (KKU), and Nalukandudde (NKD) are shown as colored circles; those with no mosquitoes collected are shown as black circles. Roads and structures are shown in black.

Mosquitoes morphologically identified as *An. gambiae s*.*l*. were individually stored in tubes with desiccant. A mosquito leg was removed for molecular species identification at the Uganda Virus Research Institute, and the remaining carcass was shipped to Notre Dame, where each mosquito was individually processed. DNA was extracted using a CTAB protocol (Chen et al. 2010), and molecularly identified to species using the SINE200 assay (Santolamazza et al. 2008). All specimens were identified as *An. gambiae s*.*s*. DNA was transferred to 96-well plates and shipped to GT-Seek LLC for library construction and sequencing (Campbell et al. 2015). Bioinformatic processing at Notre Dame was performed as described in Methods 2.1. Of the 733 mosquitoes sampled on Jaana, 716 were successfully genotyped. An additional two were excluded because metadata (sex and blood feeding status) were in conflict, resulting in 714 mosquitoes retained for CKMR analysis.

### 2.5 Latent Kinship Classification of Close-kin Pairs

Given the challenges of setting a threshold for hard-to-separate kinships, we adopted an alternative approach to pairwise close-kinship inference that makes full use of the genetic data and considers true kinship as a latent variable (Bravington et al. 2016). We performed pairwise comparisons across the observed allele frequencies on Jaana using the function ‘pairwise_kin_logl_ratios’ of the R package CKMRsim. This function calculates *L*_*ijk*_, the pairwise log-likelihood of individuals *i* and *j* for kinship category *k*. This and other parameters and symbols used in this study are summarized in Table S3. The parameter ‘numer’ of the ‘pairwise_kin_logl_ratios’ function was set to iterate over available kinship categories in the ‘kappas’ matrix of CKMRsim, including parent-offspring, full- and half-siblings, avuncular, cousins, and unrelated. The parameter ‘denom’ was set to NULL to report the pairwise log-likelihood for kinship category *k* instead of pairwise log-likelihood ratios. For each pair, we made 1,000 draws from a categorical distribution of exponentiated normalized log-likelihoods, resulting in 1,000 possible realizations of kinship pairs. For calculation of the CKMR pseudo-likelihood (Equation 4, Methods 2.6.3), we averaged over the likelihood of each realization. Further details about this approach are described in Methods 2.6.3.

### 2.6 CKMR modeling of *An. gambiae*

#### 2.6.1. Model of mosquito life history

We modeled mosquito population dynamics following Sharma et al. (2022) based on a discrete-time version of the lumped age-class model with a daily time step. However, we represented all sub-adult mosquito life stages as a single juvenile age-class because our study design was limited to adult collections. Simplified for computational purposes, the model assumes that after adult emergence, females mate immediately and only once, at random. The day after mating, she oviposits *β* eggs daily until death. The male mosquito seeks to mate every day until death, and every male mates once on average. Adult mosquito lifespan is limited to *T*_*A*_ days with an age-invariant daily mortality rate of *µ*_*A*_. Adult population abundance is constant, and spatial structure is not accounted for. Juveniles need *T*_*J*_ days to mature from egg into adult mosquitoes and are subject to a constant daily mortality probability of *µ*_*J*_. In an extension of the model of Sharma et al. (2022), we also allow for an additional source of mortality for juveniles due to the failure of a clutch to produce adult offspring, which occurs with probability *µ*_*s*_. It is important to model *µ*_*s*_ separately from *µ*_*J*_, due to the fact that siblings experience non-independent mortality risk as juveniles.

#### 2.6.2 Kinship probabilities overview

Kinship probabilities are calculated following Sharma et al. (2022) for the simplified life history stages considered here (juvenile and adult). They are based on the principle of expected relative reproductive output, where *E*_*K*_ is the total expected surviving adult offspring of a female in class *K*, relative to that of all adult female mosquitoes in the population, *E*_*A*_. The kinship probability *P*_*K*_ of two individuals *i* and *j* collected on day *t*_*i*_ and *t*_*j*_ given the kinship category *K* can be written as

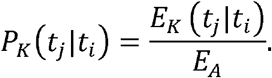

Note that absolute female abundance enters this equation implicitly in the denominator through *E*_*A*_, as elaborated in Equation (2), below.

To develop an expression for *E*_*A*_, we begin by considering the reproductive output of a single egg clutch (*R*_*c*_). An initial expression of *R*_*c*_ considers the number of surviving adult offspring on the day of sampling *t*_*2*_, after oviposition on day *y*_*2*_ as

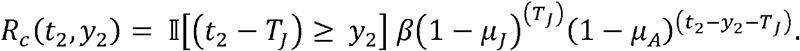

This is calculated by multiplying the number of eggs (*β*) oviposited on *y*_2_ by the daily survival rate (calculated by subtracting the daily mortality rate from one) for juveniles or adults. If the time difference between oviposition on day *y*_*2*_ and sampling day *t*_2_ is smaller than the juvenile maturation time *T*_*J*,_ then the number of surviving adult offspring is zero as indicated with the indicator function II[·].

However, this initial expression of *R*_*c*_ does not account for the phenomenon of clutch failure. Multiple studies of *An. gambiae* natural larval habitats have shown that the abundance of larvae in natural breeding sites may be decoupled from actual habitat productivity, *i*.*e*., the emergence of adult mosquitoes (Service 1973; Mutuku et al. 2006; Munga et al. 2007). Here we modify the expression of *R*_*c*_ by introducing a clutch failure rate *µ*_*S*_, allowing for some clutches to yield no adult offspring, yielding

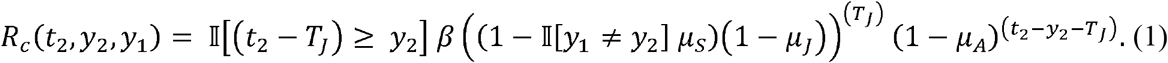

We then calculate *E*_*A*_ by summing over all oviposition events during the life of a female mosquito and multiplying by the number of adult female mosquitoes *N*_*F*_ in the population to produce

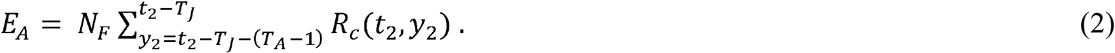

*E*_*A*_ is independent of time, because the CKMR method assumes a closed population at equilibrium with constant adult population size. We sum over the possible days of egg-laying, from the earliest (*y*_*2*_ = *t*_*2*_ - *T*_*J*_ - (*T*_*A*_-1)) to the latest (*y*_*2*_ = *t*_*2*_ - *T*_*J*_), consistent with the adult offspring being present on the day of sampling.

In Supplemental Text S1, we detail expressions for kinship probabilities *P*_*K*_ for three kinship types included in our CKMR model: mother-offspring, father-offspring, and full-siblings. The derivations reflect the fact we sampled only adults, adult sampling was lethal, and age was uncertain, but sex was known. In each case, kinship probabilities are calculated by a daily iteration over the juvenile age-class followed by the adult age-class.

#### 2.6.3 Pseudo-likelihood overview

We estimated the kinship-specific pseudo-likelihoods by summing over the binary indicator *K*_*ij*_ that a sample pair *i-j* has kinship *K*. Then, summing over the pseudo-likelihoods for mother-offspring, father-offspring, and full-sibling pairs produces the joint pseudo-likelihood,

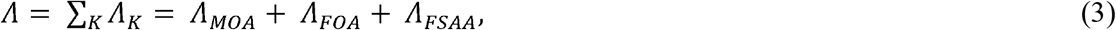

used in this CKMR analysis to make demographic inferences, which we refer to as the PO+FS CKMR model. In Supplemental Text S2, we detail expressions for *Λ*_*K*_ for these three kinship categories.

We differ somewhat from Sharma et al. (2022) in our pseudo-likelihood calculations, because those authors assumed that genetic inference of kinship was made without error. Here, we account for kinship uncertainty through latent kinship. The key issue is that inference of population size in the CKMR approach depends on the number of pairs of each kinship category, yet uncertainty about the kinship category of each pair means that there is uncertainty about the number of pairs of each kinship category. In other words, pair-level uncertainty propagates into count-level uncertainty. To address this, we made 1,000 draws of the kinship category of each pair *i,j* based on *L*_*ijk*_ from CKMRsim. Conditional on the pairwise kinships for each draw *d*, we calculated kinship-specific pseudo-likelihoods Λ_*MOA,d*_, Λ_*FOA,d*_, and Λ_*FSAA,d*_ for mother-offspring, father-offspring, and full-sibling kinships, respectively. Then, we computed the overall pseudo-likelihood

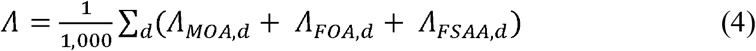

to marginalize over pair-level uncertainty about kinship type.

#### 2.6.4 Parameter estimation

We estimated four demographic parameters (*N*_*F*_, *µ*_J_, *µ*_S_, *µ*_A_) of the Jaana Island mosquito collection by maximizing the pseudo-log-likelihood of the PO+FS CKMR model (summarized in Equation 3 above; detailed in Equations S12, S15, and S17 of Supplemental Text S2) using Bayesian inference and Markov chain Monte Carlo (MCMC) sampling. We followed the same steps to estimate three demographic parameters (*N*_*F*_, *µ*_J_, *µ*_A_) using an alternative CKMR model in which the clutch failure rate parameter *µ*_*S*_ was set to zero. We refer to this as the PO+FS (*µ*_S_=0) CKMR model.

For both models, prior distributions for the parameters that we estimated were informed by available information from previous studies to the extent possible, as detailed in Supplemental Text S3 and Table S4. The posterior distributions were sampled by Markov chain Monte Carlo (MCMC) using the R package BayesianTools (Hartig et al. 2023) with the Metropolis-Hastings algorithm (Metropolis et al. 1953; Hastings 1970). For each model, we ran five Markov chains initialized with a random draw from the prior distribution, followed by a burn-in period of 1,000 samples, and, finally, a sequence of 3,000 samples with a thinning rate of 5, resulting in a total of 600 samples from the posterior distribution. These settings were informed by assessing autocorrelation and convergence of the final Markov chains with the BayesianTools and coda R packages (Plummer et al. 2006).

Fixed parameters in the model included juvenile maturation age *T*_*J*_, the number of eggs laid per daily oviposition *β*, and the maximum adult age *T*_*A*_. The juvenile maturation age *T*_*J*_ can only be inferred with parent-offspring pairs, which were not observed in our field-collected samples. We therefore assumed a maturation age of 15 days based on an average 24-h temperature on Jaana Island at 2 m above ground of ~23.6C during the period of sample collection [data obtained from the National Aeronautics and Space Administration (NASA) Langley Research Center (LaRC) Prediction of Worldwide Energy Resources (POWER) Project’s Hourly 2.4.7 version accessed on 2025/01/16]. Using this temperature mean, we exploited the juvenile development results of Bayoh and Lindsay (2003). These authors reported a temperature range between 16-34 °C where juvenile development time increased linearly for *An. gambiae* in proportion to temperature, and we extrapolated from their data to arrive at our estimate of 15 days.

#### 2.6.5 Posterior predictive checks

Posterior predictive checks on the PO+FS and PO+FS (*µ*_S_=0) CKMR models were performed to assess how well the sampled parameters were able to predict the numbers of parent-offspring pairs and full-sibling pairs actually observed in our field collected samples. We simulated numbers of each kinship category for each of 600 joint draws from the posterior distribution of the four estimated parameters, drawing the same numbers of mosquitoes as were collected each day on Jaana Island (Table S5). These simulations followed the same distributions as in the pseudo-likelihood functions (Equations S12, S15, and S17 of Supplemental Text S2).

### 2.7 CKMR model validation using individual-based simulation

To validate our estimates of demographic parameters (*N*_*F*_, *µ*_J_, *µ*_S_, *µ*_A_) on Jaana Island, we used individual-based simulations with a more realistic growth model to simulate data for validating the CKMR model (see Supplemental Text S4 for a summary of *Anopheles gambiae* life history characteristics). The SLiM version 3.7 forward-in-time genetic simulation framework (Haller and Messer 2019) with the non-Wright-Fisher option supporting overlapping generations was used for the implementation of daily mosquito maturation, emergence, and death for a single closed panmictic mosquito population at equilibrium. A detailed description of the simulation model is provided in Supplemental Text S5.

An example simulation demonstrating that the simulations exhibited their intended behavior is shown in Fig. S2. In total, we simulated 40 replicate populations ranging from ~8,000 (roughly 0.25x the estimated female population size) to ~45,000 (roughly 1.5x the estimated female population size) uniformly on a log scale. Simulation model parameters were set in such a way that they resulted in equivalent processes of mosquito emergence and mortality, despite subtle differences between the formulations of the deterministic CKMR model equations and the stochastic SLiM simulation algorithm.

## 3. Results

### 3.1. A novel *An. gambiae* genotyping panel provides power for kinship inference

We developed a 291-locus amplicon panel of microhaplotype markers distributed across all chromosome arms, excluding heterochromatic regions and common inversion polymorphisms (Fig. 2A; Table S6). Locus lengths ranged from 81 to 141 bp (mean = 125 bp). In pilot samples from island populations in Lake Victoria, the number of haplotypes per locus ranged from 3 to 32 on Bukasa and from 2 to 28 on Sserinya, with mean expected heterozygosity across loci of approximately 0.8 (Fig. 2B-C; Table S7). In contrast, the laboratory G3 colony showed markedly reduced polymorphism, with only 133 loci polymorphic and a mean expected heterozygosity of approximately 0.4 (Table S7).

**Figure 2.**
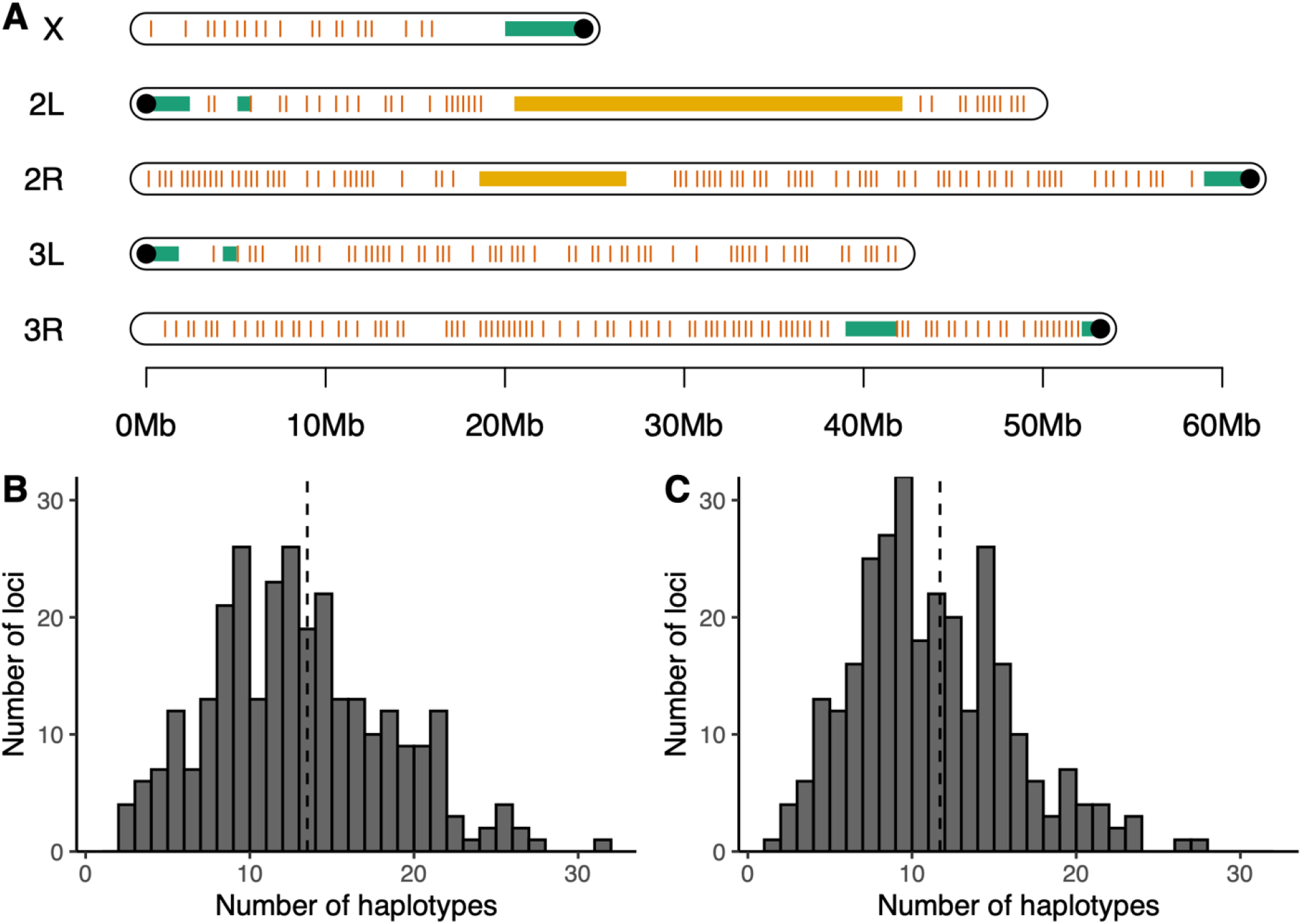
Microhaplotype markers and diversity. (A) Physical map of the 291 microhaplotype markers (red vertical lines) across the *An. gambiae* genome. Positions of inversions 2La and 2Rb are shown in orange, heterochromatic regions in green, and centromeres as black circles. (B, C) Frequency distributions of the number of microhaplotypes per locus across the 291 marker loci in *An. gambiae* samples from (B) Bukasa and (C) Sserinya. Dashed lines indicate the mean number of haplotypes per locus.

From Jaana Island (Fig. 1), 714 adult *An. gambiae* were successfully genotyped and retained for analysis. Individuals were genotyped at a mean of 260 loci (range 39-285; Fig. S3). All subsequent analyses were based on these genotype data.

### 3.2. Threshold-based kinship classification is unreliable for mosquitoes

We evaluated the performance of threshold-based kinship classification using simulated genotype pairs generated under allele frequencies from a natural island population sample and from the laboratory G3 colony. Likelihood ratio distributions were generated for parent-offspring, full-sibling, half-sibling, and unrelated kin relationships (Fig. 3; Fig. S4), and classification thresholds were selected to maximize overall accuracy (Fig. S5). False-positive and false-negative rates are summarized in Table S8.

**Figure 3.**
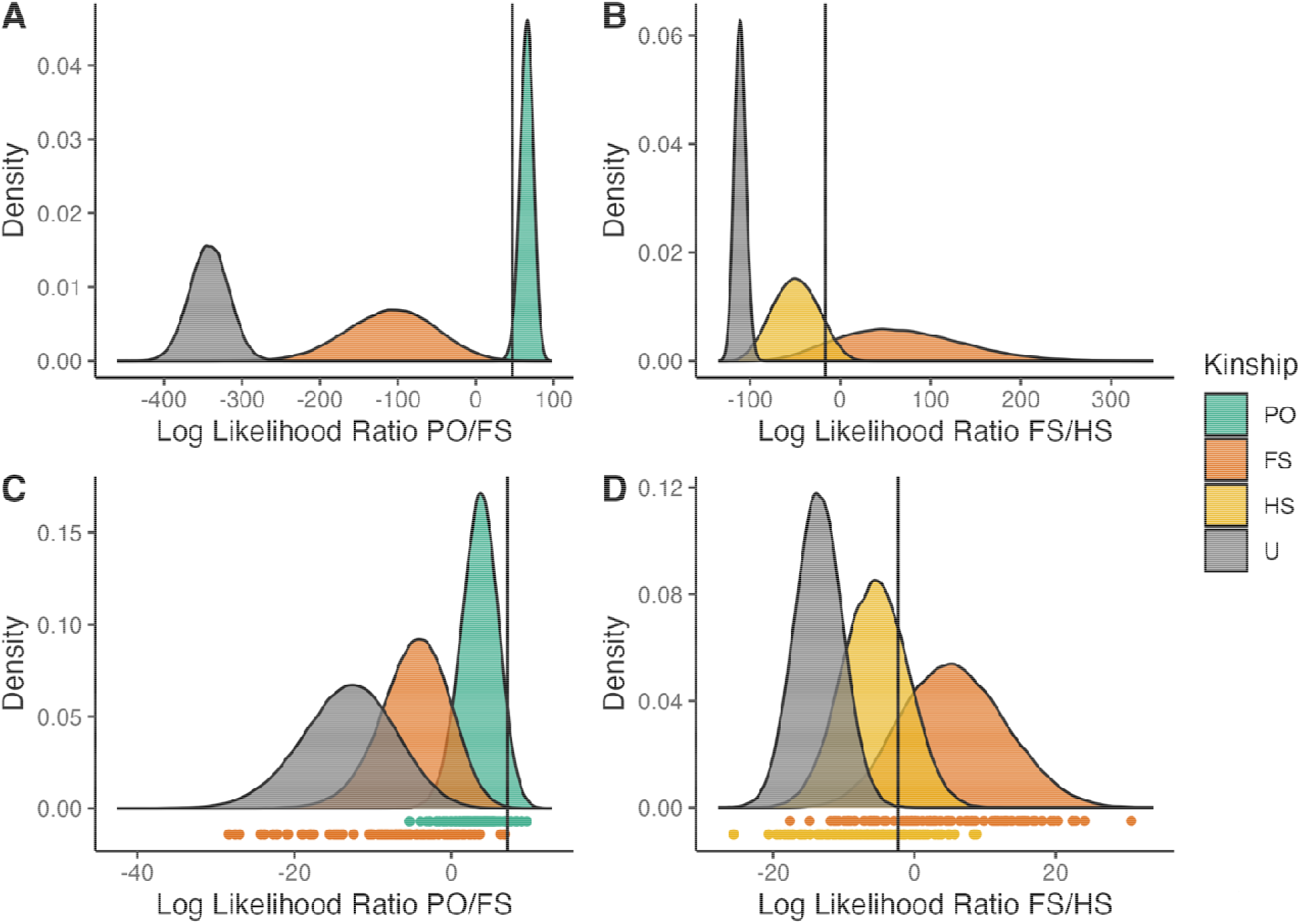
Density distributions of log-likelihood ratios for kin-pair classification. Likelihood ratios were generated from genotype pairs simulated using allele frequencies from Bukasa (A, B) or from unrelated mosquitoes in the G3 laboratory colony (C, D). In panels C and D, log-likelihood ratios computed for individual genotype pairs with known kinship from the G3 pedigree (parent–offspring, full-sibling, and half-sibling) are plotted as points beneath the density distributions, colored by kinship category. Vertical lines indicate the classification thresholds used. Abbreviations: PO, parent–offspring; FS, full-siblings; HS, half-siblings; U, unrelated.

In simulations based on G3 allele frequencies, classification accuracy was low across all kinship categories (Fig. 3C-D). In contrast, simulations based on natural population allele frequencies showed that unrelated pairs were reliably distinguished from first-degree relatives, with overall classification accuracy exceeding 0.999 (Fig. 3A-B; Table S8). However, likelihood ratio distributions for full-sibling, half-sibling, and avuncular relationships overlapped substantially (Fig. 3B). As a result, accuracy for separating these close-kin categories ranged from 0.902 to 0.905, with elevated false-positive and false-negative rates (Table S8).

### 3.3. Kin-pair detection using latent kinship reveals full-sibling but no parent-offspring pairs

We classified all 254,541 pairwise genotype comparisons among the 714 sampled mosquitoes using an alternative approach in which true kinship is treated as a latent variable (Bravington et al. 2016). No parent-offspring pairs were inferred. In contrast, a median of 63 full-sibling pairs was inferred (95% CrI: 60-65), of which 60 were high-confidence pairs with normalized pseudo-likelihoods of at least 0.98. Sibship structure supported approximate independence among inferred full-sibling pairs. We inferred a mean of 48.3 distinct sibships (range 46-53), and 41.7 of these (86%) consisted of only two siblings (Table S9).

### 3.4. Temporal and spatial clustering of full-siblings

Mosquitoes were collected on Jaana over a 20-day period. Among the 60 high-confidence full-sibling pairs, siblings were collected a mean of 3.0 days apart (95% CrI: 2.00-3.93; Fig. S6), compared to a mean of 7.1 days for randomly paired individuals (p < 0.0001). This difference indicates non-random temporal clustering of full-siblings within the sampling window.

Sampling on Jaana occurred in two main localities approximately 2,000 m apart (Fig. 1). Of the 714 mosquitoes analyzed, a total of 613 were from Lwazzi (LWZ) and 101 from the combined Kikku/Nalukandudde (KKU/NKD) settlements. Within LWZ, 36 full-sibling pairs were inferred; within KKU/NKD, 18 full-sibling pairs were inferred; six full-sibling pairs spanned these two main localities.

To assess whether the observed distribution of full-sibling pairs was consistent with a single, well-mixed population at the island scale, we performed a randomization test in which village identities of the 714 mosquitoes were randomly reassigned. For each permutation, the numbers of full-sibling pairs within LWZ, within combined KKU/NKD, and between the two were recalculated. This procedure was repeated 100,000 times to generate null expectations. Under this null model, the expected numbers of full-sibling pairs were 44 within LWZ (null interval: 38-50), 1.2 within KKU/NKD (null interval: 0-4), and 15 between villages (null interval: 9-21). Observed counts differed significantly from these expectations (LWZ: p = 0.008; KKU/NKD: p < 0.0001; between villages: p = 0.002), rejecting the null hypothesis of rapid complete mixing of individuals at this spatial scale. Nevertheless, the six pairs shared between villages indicate some movement does occur, and if we assume for simplicity that dispersal occurs before reproduction and that the proportion of immigrants in the two localities is equal, then the observed counts suggest that proportion is about 2.5% (see Supplementary Text S6 for further details).

### 3.5. Estimating demographic parameters

Because spatial structure of full-sibling pairs was detected at the island scale, CKMR inference was restricted to mosquitoes collected from LWZ. To estimate demographic parameters, we fit two CKMR models. Both models include the parent-offspring (PO) and full-sibling (FS) kinship pair probabilities, but they differ in their assumptions about *An. gambiae* reproductive biology and therefore in the parameters included. The first model estimates only three demographic parameters (adult female abundance *N*_*F*_, juvenile *μ*_*J*_ and adult *μ*_*A*_ daily mortality probabilities), as a fourth (clutch failure parameter *μ*_*S*_) was constrained to zero. We refer to this ‘PO+FS (*μ*_*S*_ = 0)’ model as the CKMR base model. In addition to these three parameters, the ‘PO+FS’ extended model also accommodates the daily possibility that an entire clutch of eggs fails to produce adult offspring—independent of daily individual juvenile mortality. Our objective was to compare the numbers of observed kin pairs in the CKMR population sample (assessed by genotyping) to the numbers expected under the alternative fitted CKMR models as a means of checking assumptions about reproductive biology (Lloyd-Jones et al. 2023).

Posterior predictive checks showed that the base model underpredicted the number of full-sibling pairs and overpredicted the number of parent-offspring pairs (Table 1; Fig. 4). By contrast, expectation under the extended model closely matched the observed numbers of full-sibling pairs and the absence of parent-offspring pairs (Table 1; Fig. 4). On this basis, we reject the base model and favor the extended CKMR model.

**Table 1.** Observed versus (/) posterior predictions of the fitted CKMR models for the number of kin pairs. Median (and 95% credible interval) for observed kin pairs; median (and 95% posterior prediction intervals) for predicted kin pairs.

| Model | Parent-offspring (PO) pairs | Full-sibling (FS) pairs |
| --- | --- | --- |
| PO+FS ($\mu_S=0$) | 0 / 11 (4-20) | 38 (36-40) / 25 (14-40) |
| PO+FS | 0 / 1 (0-5) | 38 (36-40) / 36 (21-56) |

**Figure 4.**
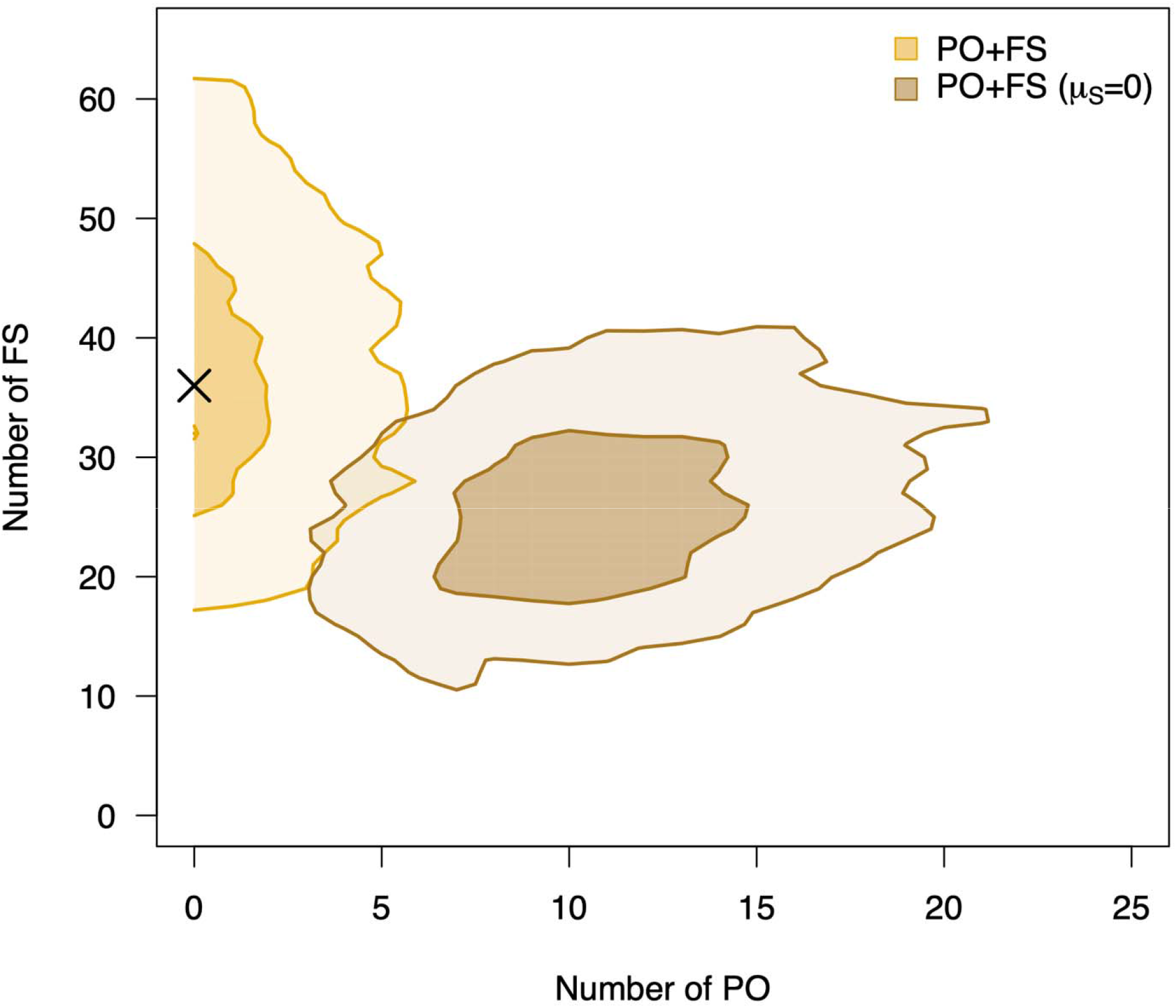
Posterior predictive checks show that modeling clutch failure is required to match observed kin-pair counts. Fitted CKMR models employed were the PO+FS (*μ*_S_=0) (brown) and the PO+FS (yellow). The solid-colored level lines represent the 50% and 95% credible intervals. The black X represents the observed number of parent-offspring (PO) and full-sibling (FS) pairs inferred from genotyping of field samples.

Adult female abundance in the extended CKMR model is estimated as 26,887 (95% CrI: 6,979 - 146,011), and daily clutch failure probability as 0.22 (0.15 - 0.31) (Table 2). Prior and posterior distributions for CKMR demographic parameters under Joint posterior samples showed a strong correlation between these two parameters (Pearson’s r = 0.97; Fig. S7).

**Table 2.** Summary of parameter estimates for the fitted extended CKMR model. Median (and 95% credibility intervals) assuming a juvenile development time *T*_*J*_ of 15 days.

| Adult female population size ($N_F$) | Juvenile mortality daily probability ($\mu_J$) | Clutch failure daily probability ($\mu_S$) | Adult mortality daily probability ($\mu_A$) |
| --- | --- | --- | --- |
| 26,887 (6,979-146,011) | 0.07 (0.02-0.23) | 0.22 (0.15-0.31) | 0.18 (0.1-0.26) |

Given the 15-day juvenile development time assumed in the model, the estimated overall probability a clutch fails is 1-(1 - 0.22)^15^ = 97.6% (95% CrI: 91.3 – 99.6%). Therefore, many females will have no surviving offspring, a few females will have many, and the effective population size will be much smaller than the census size. Using our demographic model and estimated parameter values we can calculate predicted values for the variance of female and male reproductive success of 35.0 (95% posterior predictive interval [PPI]: 12.5 – 172.6) and 41.6 (19.0 – 179.3), respectively, and an effective population size of *N*_*e*_ = 5027 (95% PPI: 2917 - 8922), giving a median ratio of effective to census size (*N*_*e*_ /(2\**N*_*F*_)) of 9.9% (2.24% – 22.6%) assuming a 1:1 adult sex ratio (see Supplementary Text S7 for details).

### 3.6 Individual-based mosquito simulations validate CKMR

To evaluate CKMR performance, true counts of parent-offspring and full-sibling pairs extracted from each simulated collection were analyzed using the extended CKMR model. Across simulated populations, CKMR estimates of adult female abundance closely matched true simulated values (Fig. 5). Counts of both parent-offspring and full-sibling pairs observed in the field data fell within the predictive intervals associated with simulated populations near the CKMR-estimated adult female abundance (Fig. 6). These results indicate that the numbers of full-sibling and parent-offspring pairs observed on Jaana are consistent with expectation for a female census size (*N*_*F*_) of ~27,000, as we estimated.

**Figure 5.**
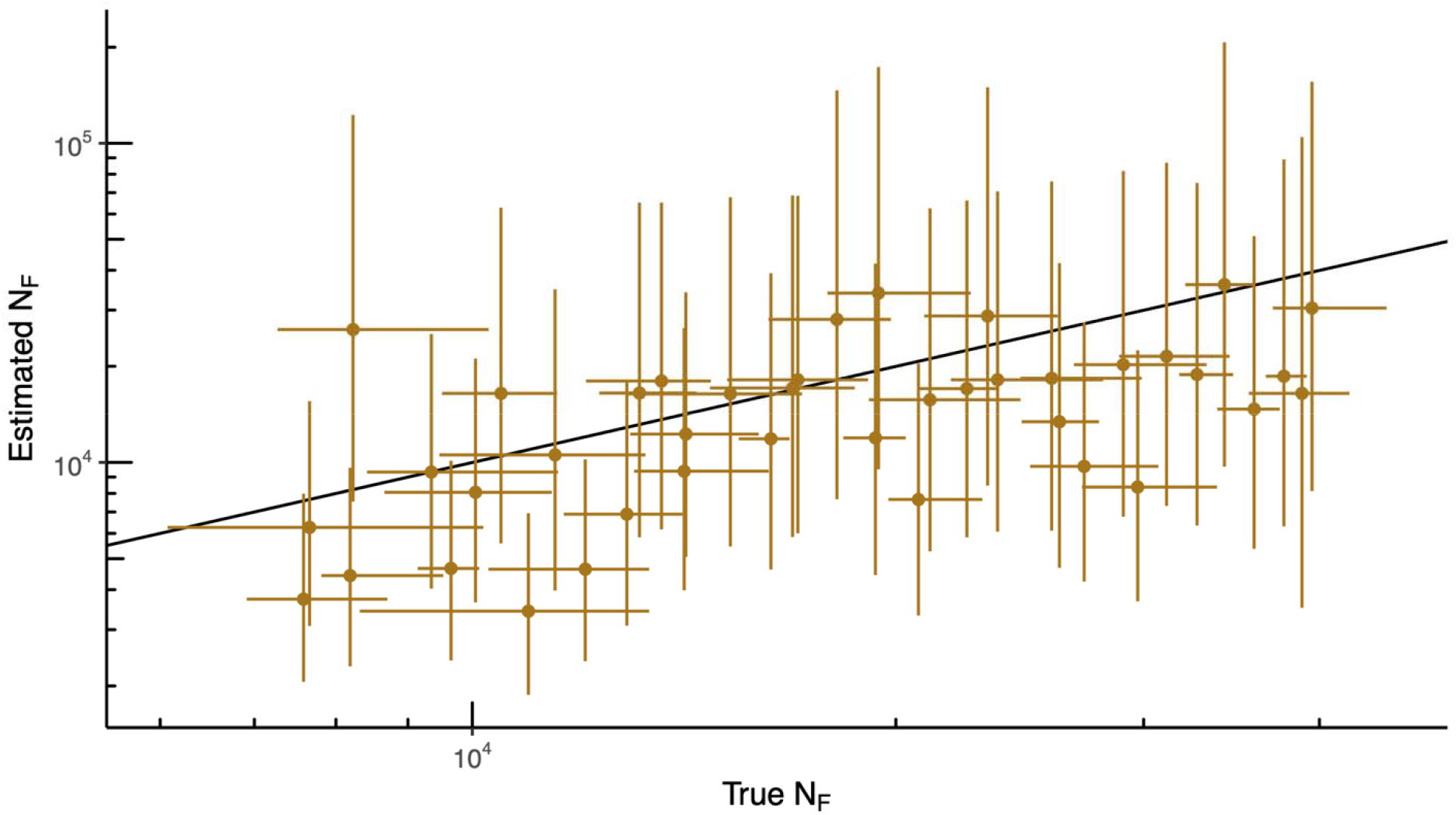
CKMR recovers adult female abundance in individual-based simulations. Median values (points), 95% credibility intervals (vertical bars), and the recorded 95% quantile intervals (horizontal bars) are shown. Credibility intervals of estimated values reflect statistical uncertainty, whereas quantile intervals of true values reflect stochastic variability. Black dotted line of slope=1 represents the expectation if true and estimated values align perfectly.

**Figure 6.**
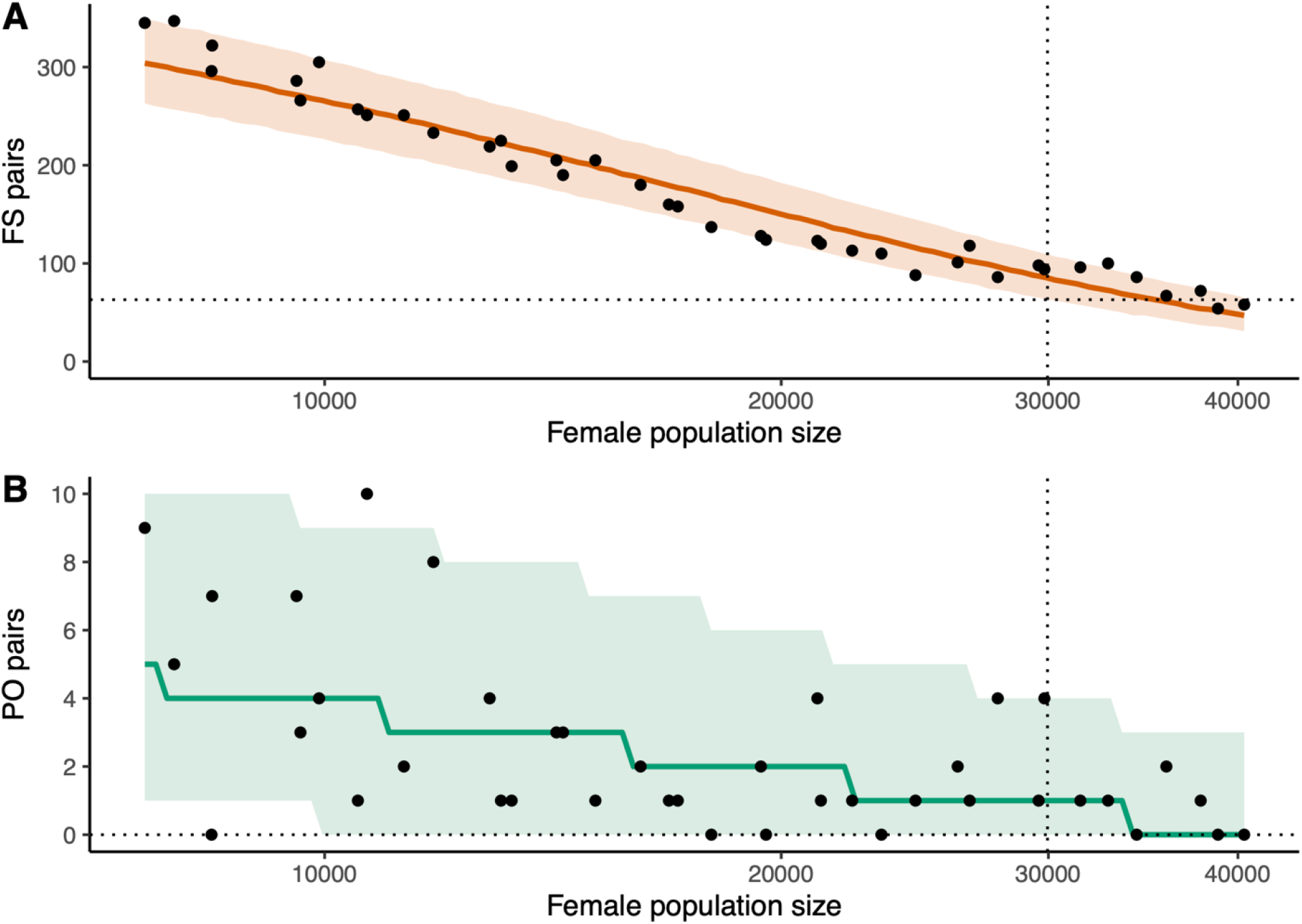
Field kin-pair counts fall within simulation-based predictive intervals near the CKMR-estimated abundance. True numbers of full-sibling (panel A) and parent-offspring (panel B) pairs extracted during simulation, shown as black circles, were fitted to a generalized linear model with a quasi-Poisson distribution. Median expected numbers (solid colored line) and the 95% predictive interval (shaded surround) from that model are shown. Black-dotted horizontal and vertical lines represent the observed number of kin pairs on Jaana Island and the female population abundance estimated by the fitted PO+FS CKMR model. Intersection of the dotted lines with the colored shading indicates a close match between estimated and true numbers. Numerical values displayed here differ slightly from those presented in Section 3.5 given that these simulations were based on parameter estimates obtained by fitting the CKMR model to data from the whole island. Accordingly, observed numbers of kin pairs presented here also reflect collections across the whole island.

## 4 Discussion

We implemented CKMR to estimate demographic parameters from a natural island population of *An. gambiae* mosquitoes. Applying a novel high-resolution amplicon sequencing panel to a sample of 714 adult mosquitoes, we detected dozens of full-sibling pairs but no parent-offspring pairs. We built a novel CKMR model that jointly analyzed the frequency of full-sibling and parent-offspring pairs and accounted for kinship uncertainty and clutch failure. We used this model to estimate adult female census size on the order of 27,000 individuals and quantify for the first time the fraction of clutches that fail. Individual-based simulations matching the empirical sampling design reproduced the observed kin-pair patterns and validated parameter recovery across a range of population sizes.

These results demonstrate that CKMR can yield informative abundance estimates in a short-lived, highly fecund insect, when demographic models explicitly accommodate mosquito life history and uncertainty in relationship inference.

### 4.1 Kin-pair distribution in *An. gambiae* affects CKMR application

As a ‘rule of thumb’ guideline for parent-offspring-only applications of CKMR, Bravington et al. (2016) suggest a sample size of 10√*N*, where *N* represents average adult abundance. Unfortunately, *An. gambiae* is generally understood to be a (seasonally) highly abundant species across its tropical African home range. *Anopheles gambiae* populations from mainland Africa are typically characterized by very high levels of nucleotide diversity and exceptionally large Ne (Miles et al. 2017). Two comparable MRR studies suggest mainland population abundance during the rainy season on the order of 150,000-300,000 females from one study (female-based MRR; Costantini et al. 1996) and 100,000-500,000 males from another (male-based MRR; Epopa et al. 2017). Several studies have shown that island populations, in particular Ssese island populations, are smaller than those on the mainland, and that connectivity with the mainland is diminished (Kayondo et al. 2005; Lukindu et al. 2018; Wiltshire et al. 2018; Bergey et al. 2020; Mwima et al. 2026). As meeting the target sample size was unfeasible for a mainland site, Jaana was chosen as the location for the first application of CKMR to a natural population of *An. gambiae* mosquitoes because—at only ~3-4 km long and ~1-2 km wide—it is one of the smallest and most isolated of the inhabited islands of the Ssese archipelago offshore Uganda. Although we indeed estimate much lower female population abundance on Jaana island (~27,000) compared to mainland MRR estimates, the absence of parent-offspring pairs initially surprised us. Even in the unlikely case that the clutch failure dynamics inferred on Jaana to explain this absence does not apply to mainland populations, their much larger size suggests that attempting a parent-offspring-only CKMR application for mainland *An. gambiae* populations might be futile. This is an important consideration, because limiting CKMR modeling only to parent-offspring pairs has been one suggested avenue to avoid the problem of concurrent and confounded close-kin relationships.

Our investigation revealed that not only do the likelihood ratio distributions of several types of kin categories overlap to an important degree (including full-siblings and half-siblings), but also our individual-based simulations demonstrate that we should expect non-trivial numbers of these difficult-to-distinguish kin types over a range of population sizes (Table S11). This is especially true for avuncular kin-pairs that are indistinguishable genetically from half-siblings using the genotyping approach taken here. For these reasons, implementation of CKMR with *An. gambiae* must rely on considering true kinship as a latent variable, to our knowledge the first application of this approach to genetic identification of kin-pairs in a CKMR framework. The traditional approach of classification with allowance for error would lead to elevated false negative and false positive error rates, biasing demographic estimates to an unacceptable degree.

Vector biologists have recognized the problem of overlapping generations for kinship prediction in extending the CKMR framework to mosquitoes. Working with a very different mosquito, the dengue vector *Aedes aegypti*, they devised a sampling design to ameliorate the problem. Deploying ovitraps only for the length of a single mosquito gonotrophic cycle, and sampling juveniles from those ovitraps, ensures that the only close-kin categories represented should be full-siblings, half-siblings, and cousins (e.g., Jasper et al. 2019; Schmidt et al. 2023). Crucially, this sampling design is highly effective for *Aedes* due to its strong preference for ovipositing in containers, typically human-made (but also tree holes) (Chadee 2004). As *An. gambiae* does not share this preference, ovitrap sampling likely would be ineffective. However, alternative CKMR sampling designs are worth exploring for *An. gambiae*.

In vertebrate applications of CKMR, age is an influential individual covariate within the models, because age at the time of sampling marks the birth year (the ‘time of marking,’ by analogy with MRR methods) (Bravington et al. 2016). Inaccurate age inference can bias parameter estimates for vertebrate species. Although knowledge of the age of individuals may not be essential for mosquito CKMR applications, it would be required to discern which individual of a parent-offspring kin pair was the parent versus the offspring (if such pairs were sampled), and it could help differentiate otherwise indistinguishable half-sibling, avuncular, and grandparent-grandchild kinships. Longstanding methods of determining age in mosquitoes include MRR and microscopic observations of morphological changes in the ovary related to the number of ovipositions (Mohammad et al. 2024, and refs. therein); both methods are labor-intensive, lack precision, and are impractical to apply in conjunction with CKMR sampling. The development of newer approaches is being driven by the epidemiological relevance of predicting mosquito age, given the positive relationship between age and vector-borne disease transmission. Candidate tools include transcriptional profiling and spectroscopic techniques (Mohammad et al. 2024, and refs. therein), but cost and sensitivity remain challenges; none have seen widespread field adoption to date.

### 4.2 Demographic estimates are female-biased

We report adult daily survival probabilities and population abundance estimates only for females. Males comprise a relatively small fraction (14%) of the 714 mosquitoes collected and analyzed, an unsurprising outcome given the indoor-resting adult mosquito sampling performed. Although a 1:1 sex ratio is expected at the juvenile stage, evidence suggests that adult *An. gambiae* males have shorter lifespans than females (Lambert et al. 2022), and might represent a smaller fraction of the adult population. Future CKMR applications to this species should attempt to collect more males, ideally through sampling of male swarms if they can be located, and analyses should model the sexes independently. For the present, we urge caution in assuming that total population density is double that estimated for adult females.

### 4.3 Interpreting the absence of parent–offspring pairs

At face value, the absence of parent-offspring pairs might suggest a very large population, while the abundance of full-sibling pairs might suggest a small population. However, these conflicting interpretations implicitly assume relatively homogeneous reproductive success, an assumption violated in *An. gambiae*. In this system, many oviposition events fail entirely for a variety of reasons: small larval habitats are ephemeral and frequently destroyed by flooding or evaporation, or juveniles succumb *en masse* to predation, parasitism or nutritional deficits. As a result, recruitment within a short sampling window is dominated by offspring from a small number of successful clutches, inflating sibling recovery while suppressing parent-offspring detections.

The importance of clutch failure in *An. gambiae* population dynamics is validated by comparing observed versus expected kin under the base and extended CKMR models. Under the base model, daily juvenile mortality estimates are unrealistically extreme (compare Table S3 to Table 2), and no combination of parameter values reproduces the observed PO and FS pair numbers (Fig. 4; Table 1). Although future applications of CKMR to *An. gambiae* will benefit from careful consideration of minimum sampling requirements, we can reject an alternative explanation for the lack of observed PO pairs that faults the sampling design (*e*.*g*., the short sampling window and exclusive reliance on indoor-resting adult collections). Our posterior predictive checks simulated new data under both fitted CKMR models using the same sampling duration and same number, sex, gonotrophic stage, and sampling day to match empirical mosquito collections. Under the base model lacking the breeding-site failure parameter, 95% of those simulations led to an expected number of PO pairs between 4 and 20 (Table 1).

The strong posterior correlation between adult female abundance and clutch failure reflects partial non-identifiability between these parameters, a feature expected in systems characterized by sweepstakes-like reproduction, in which a small fraction of reproductive events contributes disproportionately to recruitment (Hedgecock and Pudovkin 2011; Palstra and Fraser 2012). Nevertheless, posterior predictive checks and individual-based simulations show that only models allowing for clutch failure reproduce both the observed number of full-sibling pairs and the absence of parent–offspring pairs. In this system, the absence of parent–offspring pairs is therefore not reflective of population size alone, but instead must be interpreted in the context of reproductive variance.

A consequence of the high probability of total clutch failure is that the variance of reproductive success is predicted to be much larger than the default Poisson assumption in many population genetic models, and the effective population size much smaller than the census size. Such a disparity between effective and census size is expected in systems characterized by sweepstakes-like reproduction (Hedgecock and Pudovkin 2011; Palstra and Fraser 2012). A recent synthesis advocated joint evaluation of the ecological and evolutionary consequences of abundance (Waples and Feutry 2022). In that vein, Babyn et al. (2024) described a method to estimate variances in reproductive success and effective population size from CKMR data in vertebrate populations, by comparing sibship patterns within and between annual cohorts. With the same objective, but using *An. gambiae* data from our CKMR demographic model and a different analytical approach, we predict high values for the variance in reproductive success consistent with sweepstakes reproduction, and estimate an effective population size of approximately 5,000. That estimate agrees well with previous genomic estimates of effective population size in the Lake Victoria region, which report *N*_*e*_ values orders of magnitude lower than plausible census sizes (Wiltshire et al. 2018; Mwima et al. 2026).

### 4.4 Future prospects for genetic control programs

Estimates of adult female abundance are central to the design, implementation and evaluation of genetic mosquito control strategies, including sterile male release, *Wolbachia*-based interventions, and gene drive systems aimed at population suppression. Beyond abundance estimation, the spatial distribution of close-kin pairs can provide information relevant to dispersal and population structure at scales important for control operations. In our dataset, the presence of full-sibling pairs spanning villages separated by approximately 2 km indicates movement across spatial scales relevant to intervention design. Recent extensions of CKMR explicitly incorporate spatial information to jointly infer dispersal and abundance from the locations of related individuals (Jasper et al. 2022; Marshall et al. 2025). Simulation-assisted approaches, such as CKMRnn, further integrate spatially explicit demographic simulations with machine learning to recover abundance under heterogeneous or incomplete sampling designs (Patterson et al. 2026).

Together, these approaches suggest that CKMR can contribute not only estimates of adult abundance but also operationally relevant insights into dispersal and connectivity, if sampling design and demographic modeling are tailored to the biological and spatial context of the target population.

## Conclusions

Close-kin mark-recapture is a promising approach to demographic estimation in a wide spectrum of taxa, but exploration of its usefulness outside of a handful of vertebrate species has barely begun. Here, we show that CKMR can be applied successfully to a natural mosquito population when demographic models account for life-history processes that shape kinship patterns. By integrating high-resolution microhaplotype genotyping with a latent-kinship CKMR framework and explicit modeling of clutch failure, we obtained stable estimates of adult female census size that are consistent with both observed kin-pair patterns and individual-based simulations. Our results demonstrate that jointly analyzing the frequency of different classes of relatives, in particular parent-offspring and full-sibling pairs, can give refined estimates of census size and reveal signals of sweepstakes-like reproduction.

Together, these findings establish CKMR as a practical and biologically realistic approach for estimating adult mosquito abundance without the need for release experiments. More broadly, they highlight the importance of aligning demographic inference methods with species-specific life history, particularly in short-lived, highly fecund organisms. With further development to incorporate spatial structure and seasonal dynamics, CKMR has strong potential to contribute to demographic inference in vector ecology and to support the design and evaluation of mosquito control strategies.

## Acknowledgments

We thank the UVRI laboratory staff and field entomology team, and the communities of Jaana Island who consented to mosquito collection in their villages and homes. We thank Nate Campbell (GTSeek LLC) for assistance with design of the genotyping panel, Mark Benedict for assistance with the G3 controlled crosses, and Matthew Hahn, Dan Neafsey, and Christina Bergey for helpful discussion. This study was supported by a grant from the Gates Foundation and Coefficient Giving to A.B. (PI), J.K.K. and N.J.B.

## Disclaimers

The findings and conclusions in this report are those of the authors and do not necessarily represent the official position of the CDC nor the CDC Foundation. The use of trade names and commercial sources are for identification only and do not imply endorsement by the Centers for Disease Control and Prevention (CDC), the Public Health Service, or the U.S. Department of Health and Human Services. Product names, logos, brands, and other trademarks featured or referred to within are the property of their respective trademark holders. These trademark holders are not affiliated with CDC. CDC does not sponsor or endorse these materials but references them as examples of materials to use for learning activities and demonstrations.

## Data Archiving Statement

All analysis pipelines are open source and can be found at https://github.com/lerch-a/Anopheles_abundance. Raw sequences (fastq files) have been deposited with NCBI SRA under bioproject number xxxxxxx and final microhaplotypes are deposited in the Zenodo repository available at DOI:10.5281/zenodo.18928671 (*to be completed after manuscript is accepted for publication*).

## Author Contributions

NJB conceived the research project with input from STS; KB and ML designed the CKMR sampling supervised by JKK; PB conducted the controlled laboratory crosses guided by EMD; STS designed the genotyping panel and the computational pipeline for bioinformatic processing of amplicon sequences hosted by NJB; AL performed the CKMR modeling, individual-based simulations, and figure production mentored by TAP; TJH modeled migration, reproductive variance, and effective population size mentored by AB; NJB, AL, STS, and TAP drafted the manuscript; AL, STS, TJH, AB, TAP and NJB edited the manuscript; all authors approved the manuscript.

## Appendices

Appendix 1. Agam_CKMR_mapped_amplicons.354

Appendix 2. Agam.amplicons.final_list

## Supplementary Material

Tables S1-S11

Figures S1-S7

Supplemental Text S1-S7

