## Supplementary Material for "Estimation of adult census size from close-kin dyads in the malaria mosquito *Anopheles gambiae*"

Supplementary Tables S1-S11

Supplementary Figures S1-S7

Supplementary Text S1-S7

References

**Supplementary Tables**

**Table S1.** ‘Kappas’ Matrix of Cotterman coefficients for select pairwise kinship relationships. Kappa 0, 1, and 2 represent the probabilities that two non-inbred individuals share zero, one, or two alleles, respectively, identical-by-descent (IBD) at a given autosomal locus.

|  | kappa0 | kappa1 | kappa2 |
| --- | --- | --- | --- |
| Parent-offspring (PO) | 0.000 | 1.000 | 0.000 |
| Full sibling (FS) | 0.250 | 0.500 | 0.250 |
| Half-sibling (HS) | 0.500 | 0.500 | 0.000 |
| Grandparent-grandchild (G) | 0.500 | 0.500 | 0.000 |
| Avuncular^1^ (A) | 0.500 | 0.500 | 0.000 |
| Double first cousin (DFC) | 0.563 | 0.375 | 0.063 |
| Half avuncular^1^ (HAN) | 0.750 | 0.250 | 0.000 |
| Full cousin (FC) | 0.750 | 0.250 | 0.000 |
| Half first cousin (HFC) | 0.875 | 0.125 | 0.000 |
| Unrelated^2^ (U) | 1.000 | 0.000 | 0.000 |

^1^Avuncular: Aunt/Uncle-Niece/Nephew.

^2^In practice, ‘Unrelated’ can also capture all kinship categories not explicitly modeled in a CKMR study

**Table S2.** *An. gambiae* sampling results for Jaana Island, Oct-Nov 2021

| Village | No. *An. gambiae* | % of total |
| --- | --- | --- |
| LWZ | 630 | 86% |
| KKU | 86 | 12% |
| NKD | 17 | 2% |
| Total | 733 | 100% |

**Table S3.** Overview of symbols and parameters used in this study.

| Symbol | Description |
| --- | --- |
| *i, j* | Individuals *i* and *j* |
| *k* | Kinship category *k* |
| *L_ijk_* | Pairwise log-likelihood of individuals *i* and *j* for kinship category *k* |
| *T_J_* | Juvenile maturation time from egg to adult |
| *T_A_* | Maximum adult age |
| *N_F_* | Adult female population size |
| *β* | Number of eggs per clutch |
| $\boldsymbol{\mu}_{\boldsymbol{J}}$ | Juvenile mortality probability per day |
| $\boldsymbol{\mu}_{\boldsymbol{A}}$ | Adult mortality probability per day |
| $\boldsymbol{\mu}_{\boldsymbol{S}}$ | Clutch failure probability per day |
| *t_i_, t_j,_ t_1_, t_2_* | Time of collection of individual *i* and *j* |
| *R_c_* | Number of surviving offspring of a single egg clutch |
| *y_1_, y_2_* | Day of oviposition relative to emergence of individual 1 and 2 |
| *y_m_* | Day of mating |
| *E_MOA_* | Expected number of surviving adult offspring for kinship category mother-offspring |
| *P_MOA_* | Probability that offspring is collected at day t_2_ given the mother was collected at t_1_ |
| *p_MOA_* | Probability that an adult sampled on day *t_2_* has a mother sampled on day *t_1_* |
| *k_MOA_* (t_2_\|t_1_) | Observed number of mother-offspring pairs with mothers sampled on t_1_ and offspring sampled on t_2_ |
| *p_A_*(t) | Probability that a given adult in the population survived to age *t.* |
| *Λ* | Joint pseudo loglikelihood |
| $\boldsymbol{\Lambda}_{\boldsymbol{MOA}}$ | Mother-offspring pseudo loglikelihood |
| $\boldsymbol{\Lambda}_{\boldsymbol{FOA}}$ | Father-offspring pseudo loglikelihood |
| $\boldsymbol{\Lambda}_{\boldsymbol{FSAA}}$ | Full-sibling pseudo loglikelihood |
| n_F_(*t_1_*) | Number of sampled adult females on day *t_1_* |
| n_M_(*t_1_*) | Number of adult males on day *t_1_* |
| n_A_(*t_2_*) | Number of sampled adults on day *t_2_* |
| n_D_ | Number of parameter draws |
| d | Parameter draw index |

**Table S4.** CKMR parameter prior distributions (median and 95% uncertainty intervals).

| Model | Adult female population size *(N_F_*) | Juvenile mortality daily probability  (𝜇_J_) | Clutch failure daily probability  (𝜇_S_) | Adult mortality daily probability (𝜇_A_) |
| --- | --- | --- | --- | --- |
| PO+FS | 4528 (252 - 77971) | 0.09 (0.02 - 0.39) | 0.21 (0.05 - 0.54) | 0.09 (0.03 - 0.28) |
| PO+FS (𝜇_S_=0) | 4491 (262 - 74343) | 0.33 (0.29 - 0.37) | 0 | 0.1 (0.03 - 0.28) |

**Table S5.** Simulated sampling scheme emulating adult mosquito collection on Jaana

| Day | 1 | 2 | 3 | 4 | 5 | 6 | 7 | 8 | 9 | 10 | 11 | 12 | 13 | 14 | 15 | 16 | 17 | 18 | 19 | 20 |
| --- | --- | --- | --- | --- | --- | --- | --- | --- | --- | --- | --- | --- | --- | --- | --- | --- | --- | --- | --- | --- |
| Female (+)^1^ | 23 | 89 | 0 | 30 | 31 | 1 | 15 | 55 | 0 | 31 | 7 | 15 | 0 | 1 | 3 | 37 | 19 | 10 | 28 | 15 |
| Female (-)^1^ | 23 | 34 | 0 | 8 | 4 | 3 | 5 | 35 | 0 | 7 | 4 | 8 | 0 | 2 | 5 | 27 | 11 | 3 | 11 | 12 |
| Male | 7 | 37 | 0 | 5 | 4 | 2 | 0 | 24 | 0 | 3 | 2 | 0 | 0 | 0 | 0 | 2 | 3 | 4 | 6 | 3 |
| Total | 53 | 160 | 0 | 43 | 39 | 6 | 20 | 114 | 0 | 41 | 13 | 23 | 0 | 3 | 8 | 66 | 33 | 17 | 45 | 30 |

**^1^**Females captured with (+) or without (-) bloodmeals

**Table S6.** Genomic distribution of 291 microhaplotype markers of the genotyping panel

| Chr Arm | No. markers | Mean Inter-Locus Distance* |
| --- | --- | --- |
| X | 20 | ~0.953Mb (range 0.266-4.080Mb) |
| 2L | 33 | ~0.683Mb (range 0.041-1.861Mb) |
| 2R | 87 | ~0.571Mb (range 0.135-2.722Mb) |
| 3L | 59 | ~0.645Mb (range 0.069-1.966Mb) |
| 3R | 92 | ~0.524Mb (range 0.009-2.413Mb) |

*excluding inverted regions on chr 2 and heterochromatic regions

**Table S7.** Genetic diversity estimated from the genotyping panel in population samples

| Samples | No. mosquitoes | Polymorphic loci | Invariant loci | Average H_E_ |
| --- | --- | --- | --- | --- |
| Sserinya | 43 | 291 | 0 | 0.785 |
| Bukasa | 51 | 291 | 0 | 0.813 |
| G3 colony | 31* | 133 | 158 | 0.405 |
| Jaana | 714 | 291 | 0 | 0.770 |

*Unrelated G3 mosquitoes

**Table S8.** Log-likelihood-ratio classification threshold and associated statistics from natural populations (Bukasa, Sserinya) or laboratory culture (G3 colony). PO, Parent-offspring; FS, Full-sibling; HS, Half-sibling; U, Unrelated.

| Test | False Negative Rate | False Positive Rate | Accuracy | Log-likelihood cut-off |
| --- | --- | --- | --- | --- |
| Bukasa |  |  |  |  |
| PO/FS | 0.02160 | 0.00046 | 0.999 | 47.2 |
| FS/HS | 0.09410 | 0.09820 | 0.905 | -16.4 |
| FS/U | 0.00051 | 0.00000 | 1.000 | -99.9 |
| PO/U | 0.00000 | 0.00000 | 1.000 | 194 |
| Sserinya |  |  |  |  |
| PO/FS | 0.01510 | 0.00074 | 0.999 | 39.2 |
| FS/HS | 0.09900 | 0.09510 | 0.902 | -14.2 |
| FS/U | 0.00101 | 0.00000 | 1.000 | -82.3 |
| PO/U | 0.00000 | 0.00000 | 1.000 | 165 |
| G3 colony |  |  |  |  |
| PO/FS | 0.94100 | 0.00061 | 0.986 | 7.1 |
| FS/HS | 0.13600 | 0.24100 | 0.829 | -2.3 |
| FS/U | 0.16600 | 0.00001 | 1.000 | 5.7 |
| PO/U | 0.03100 | 0.00000 | 1.000 | 8.9 |

**Table S9.** The number of full-sibling sibships predicted on Jaana (mean and range) sorted by sibship size.

| Sibship Size | 2 | 3 | 4 | 5 | 6 |
| --- | --- | --- | --- | --- | --- |
| No. Sibships | 41.7 (38-47) | 5.1 (3-8) | 0.6 (0-3) | 1 (1-2) | 0 (0-1) |

**Table S10. Summary of parameter estimates for the fitted** PO+FS (𝜇_S_=0) **model. Median (and 95% credibility intervals) assuming a juvenile development time *T_J_* of 15 days.**

| Adult female population size *(N_F_*) | Juvenile mortality daily probability (𝜇_J_) | Clutch failure daily probability (𝜇_S_) | Adult mortality daily probability (𝜇_A_) |
| --- | --- | --- | --- |
| 2,028 (1,412 - 3,145) | 0.29 (0.28 - 0.3) | 0 | 0.26 (0.18 - 0.35) |

**Table S11.** Expected numbers of difficult-to-distinguish kin-pairs (FS, HS, A) as female abundance changes. For reference, see Table S1.

| *N_F_* | Mother-offspring (MO) | Father-offspring (FO) | Full-sibling (FS) | Half-sibling (HS) | Avuncular (A) |
| --- | --- | --- | --- | --- | --- |
| 10,000 | 3.8 | 0.3 | 265.8 | 7.1 | 120.6 |
| 20,000 | 1.8 | 0.2 | 150.6 | 4.0 | 71.5 |
| 30,000 | 0.9 | 0.2 | 85.3 | 2.3 | 42.3 |
| 40,000 | 0.4 | 0.1 | 48.3 | 1.3 | 25.1 |

**Supplementary Figures**

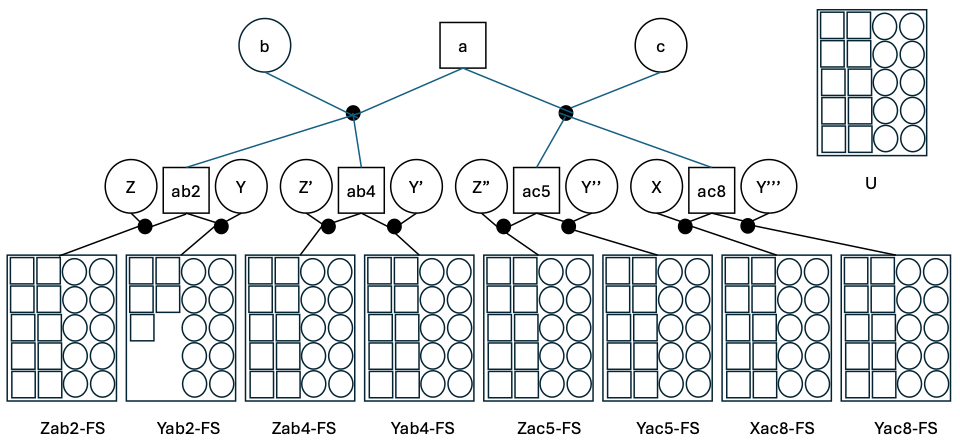

**Figure S1. Pedigree resulting from the controlled *An. gambiae* G3 crossing scheme**. Individual males (represented as squares) were mated to two females (circles) in the first and second generations (first and second rows). Eight full-sibling (FS) families were obtained in generation 3 (row 3). Comparing pairs from different families results in more distant kinship relationships (*e.g*., half siblings, cousins) relative to the closest, dominant [sensu Anderson (2022)] full-sibling relationships within a family, which was our focus here. Box at upper right labeled “U” represents twenty unrelated males and females from the G3 colony that were genotyped.

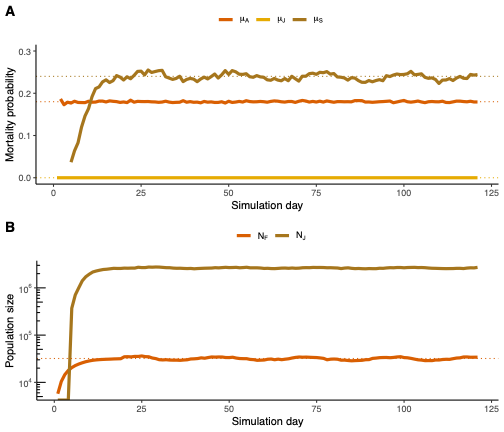

**Figure S2.** **Demographic outputs of the simulation model.** To confirm that the simulation model behaved as intended, we recorded the daily adult mortality probability (A) and population size (B) from a single realization of the individual-based simulation model, shown here with solid lines. These values closely agree with the values of model parameters (dotted lines) estimated by the CKMR model.

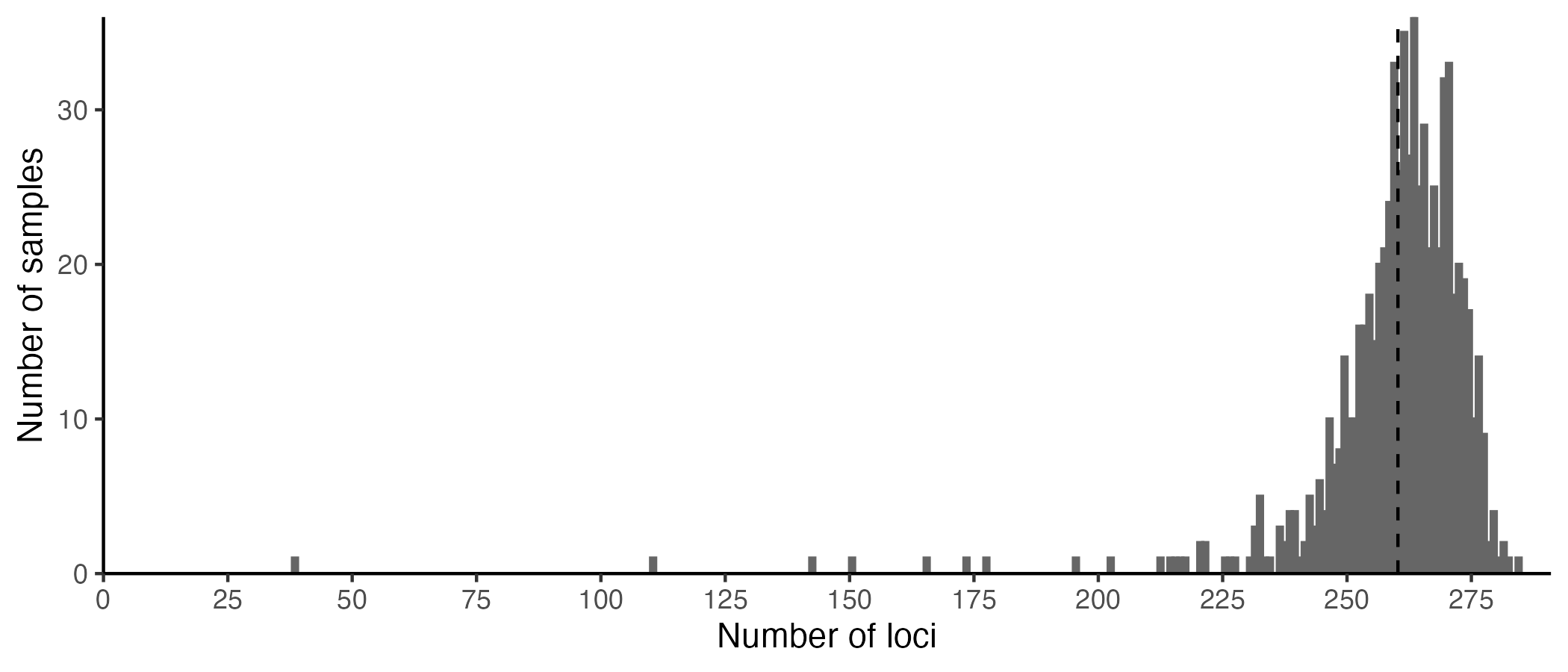

**Figure S3. Frequency distribution of the number of loci successfully genotyped per adult mosquito collected on Jaana Island**. Vertical dashed line represents the mean number of successfully genotyped loci per mosquito.

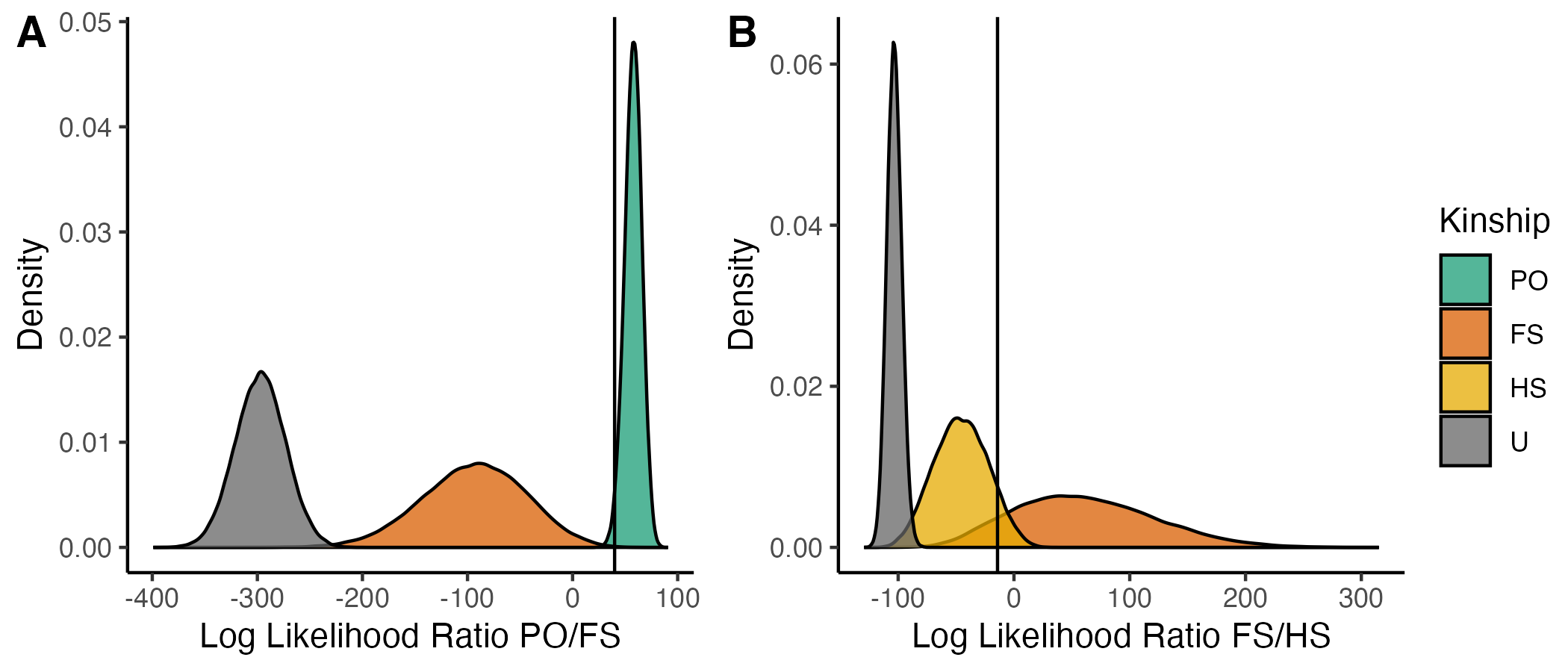

**Figure S4. Density distributions of log-likelihood ratios for kinship classification.** Likelihood ratios were generated from pairs of genotypes simulated from allele frequencies in the data from Sserinya. (A) PO/FS; (B) FS/HS. Vertical lines represent the decision-making thresholds used. PO, parent-offspring; HS, half-siblings; FS, full-siblings; U, unrelated.

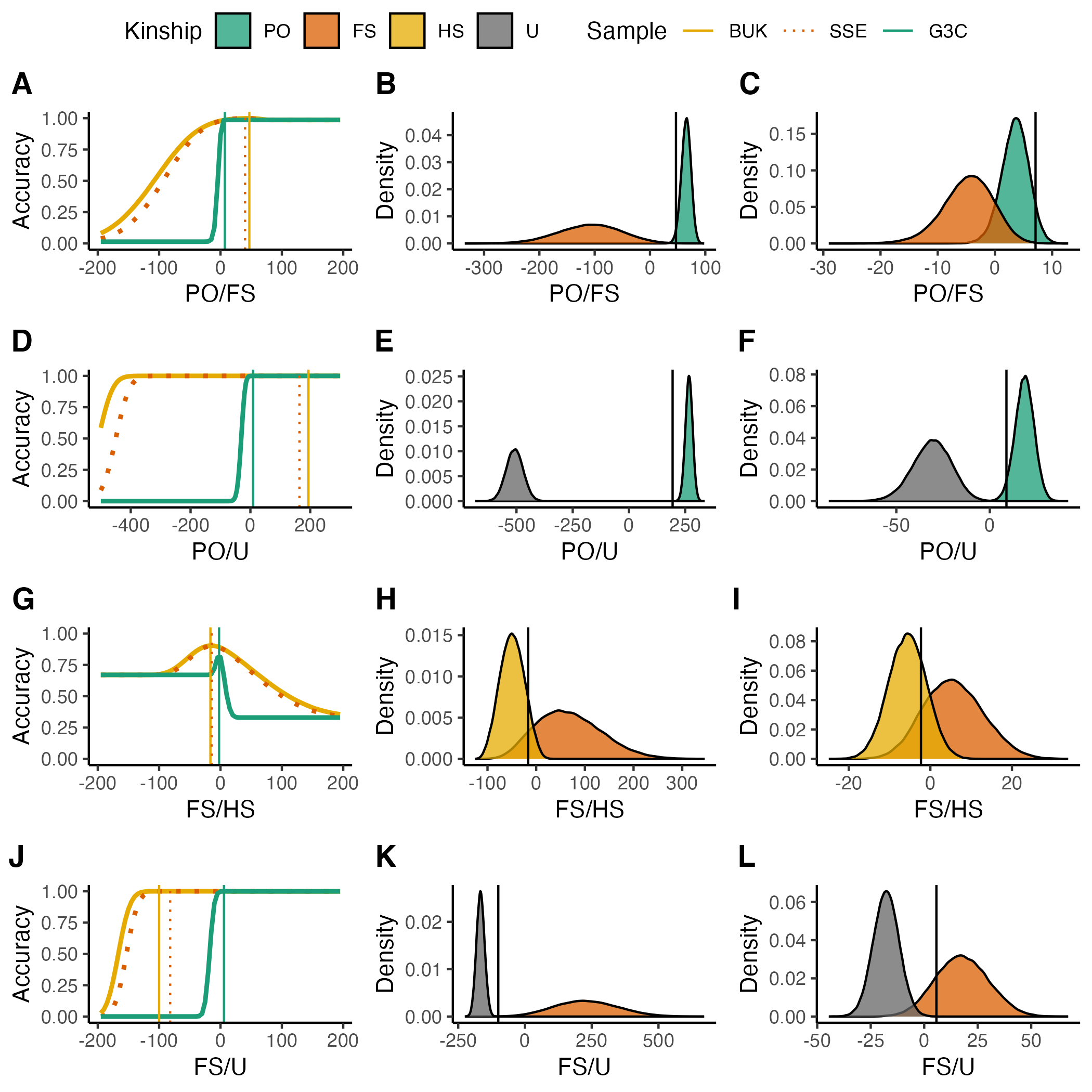

**Figure S5.** **Kinship classification accuracy as a function of threshold values of log-likelihood ratios.** For the classification-with-error approach to kinship classification, we used overall classification accuracy (y-axis in left column) as the basis for selecting threshold values of the log-likelihood ratio. Lines in the left column panels show what overall classification accuracy would be for all possible threshold values (x-axis). Threshold values that maximize overall classification accuracy are indicated with vertical lines. For reference, distributions of log-likelihood ratios among distinct pairs in each of the Bukasa (middle column) and G3 colony (right column) populations are shown. Results for different kinship categories are shown separately in each row. Parent-offspring, PO; full-sibling, FS; half-sibling, HS; unrelated, U.

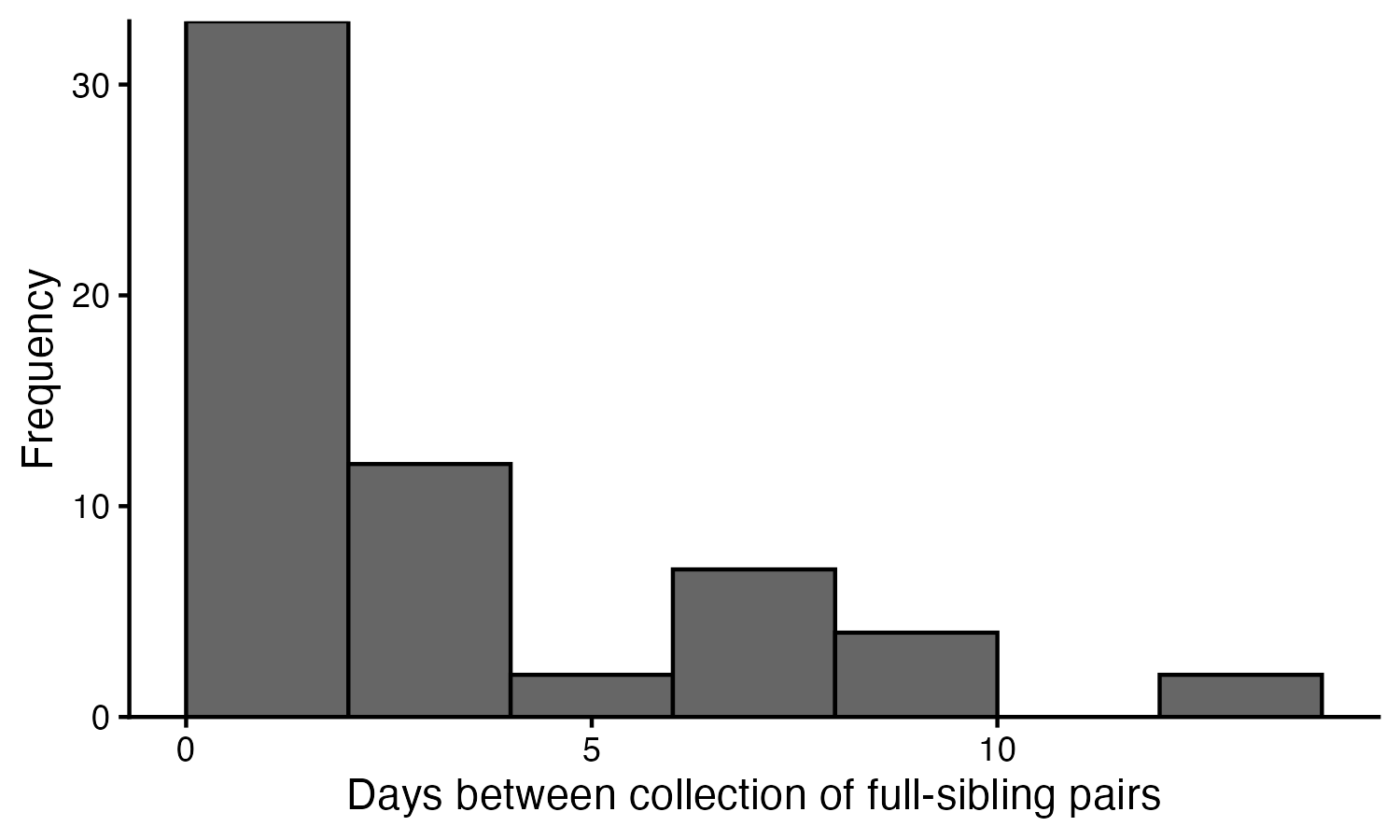

**Figure S6. Number of days between collection of full-sibling pairs.** The mean was 3.0 (95% CrI: 2.00-3.93) days.

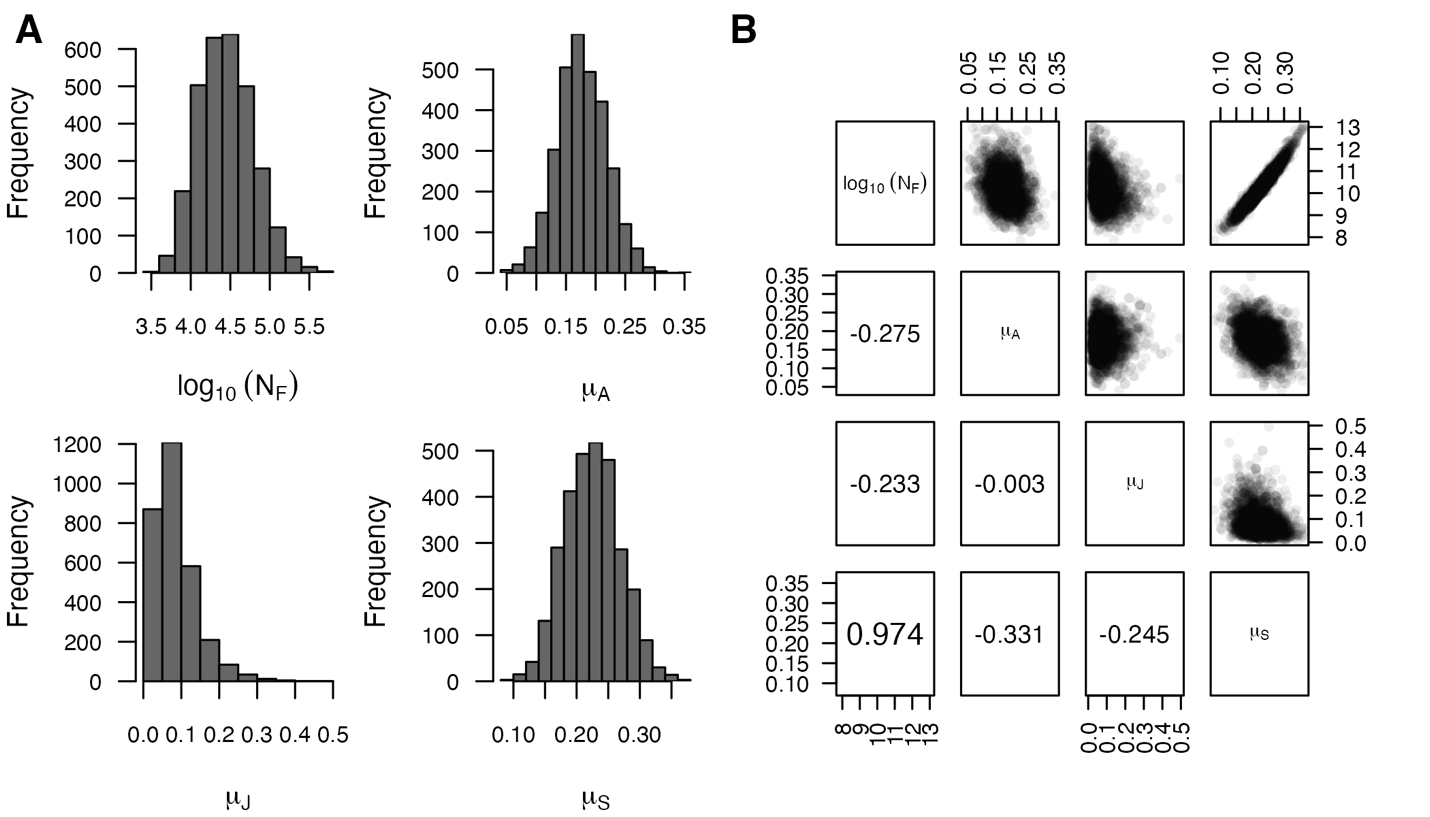

**Figure S7. Posterior distributions of estimated parameters. (**A). Marginal posterior distributions for the four parameters. (B) Above diagonal, scatterplots of the joint posterior estimates for all pairs of parameters. These plots show which parameters have estimates that are effectively independent of one another (low correlation) or that co-vary with one another (higher correlation). Below diagonal, pairwise Pearson correlations for the four parameters.

**Supplemental Text S1. Type-specific kinship probabilities.**

**Mother-adult offspring (MOA) kinship probability.** To calculate the expected number of surviving offspring on sampling day *t_2_* of a single adult female mosquito (mother) collected on day *t_1_*, we sum over all possible oviposition events *y_2_* during her lifespan (which concludes on the day of her sampling), as parity and age of collected mosquitoes are unknown. Mathematically, this is expressed by

$E_{MOA}\left( t_{2}|t_{1} \right)=\sum_{y_{2}=t_{2}-T_{j}-\left( T_{A}-1 \right)}^{t_{2}-T_{J}} \left( 1-\mu_{A} \right)^{\left( t_{1}-y_{2} \right)}\left( \mathbb{I}\left[ \left( t_{1}-{(T}_{A}-1) \right) \leq y_{2}\leq t_{1} \right] \right) R_{c}\left( t_{2}, y_{2} \right)$. (S1)

Owing to a daily mortality risk, older mothers are less likely, thus each oviposition event *R_c_* is multiplied by the survival probability of the female mosquito mother at oviposition.

Given an adult female collected on day *t_1_*, the probability that an adult mosquito collected on day *t_2_* is her offspring is

$P_{MOA}\left( t_{2}|t_{1} \right)=\frac{E_{MOA}\left( t_{2}|t_{1} \right)}{E_{A}}$.	(S2)

**Father-adult offspring (FOA) kinship probability.** To estimate the number of surviving offspring on day *t_2_* of a single adult male mosquito (father) collected on day *t_1_*, we sum over all possible mating events during the lifespan of the father mosquito (Equation S3) and all oviposition events during the lifespan of the mated female mosquito (Equation S5). The earliest mating event *y_m_* is the day after adult emergence and the latest is at the end of his lifespan at sampling. Each potential mating event is multiplied by the survival probability *p_A_* of the male mosquito (father), which yields

$E_{FOA}\left( t_{2}|t_{1} \right)=\sum_{y_{m}=t_{1}-\left( T_{A}-1 \right)}^{t_{1}} p_{A}\left( t_{1}-y_{m} \right)E_{FOA}\left( t_{2}|t_{1,}y_{m} \right)$.	(S3)

The term *p_A_(t)* represents the probability that a given adult in the population survived to age *t*, and following from the daily adult survival probability (1 − *μ_A_*), is given by

$p_{A}\left( t \right)=\frac{\left( 1-\mu_{A} \right)^{t}}{\sum_{y_{m}=0}^{T_{A}-1} \left( 1-\mu_{A} \right)^{y_{m}}}.$	(S4)

Under the simplified model assumed by Sharma et al. (2022), the mother mosquito can lay eggs from the day of mating (*y_2_=y_m_*) until she dies (*y_2_=y_m_ + (T_A_-1)*). The possible oviposition events from the perspective of the offspring collected on day *t*_2_ limits the timespan further, from *y_2_* = *t_2_* - *T_J_* - *T_A_* to *y_2_* = *t_2_* - *T_J_* . To account for mortality risk, each oviposition event *R_c_* is multiplied by the survival probability of the female mosquito (mother) at oviposition, resulting in

$E_{FOA}\left( t_{2}|t_{1,}y_{m} \right)=\sum_{y_{2}=y_{m}}^{y_{m}+\left( T_{A}-1 \right)} \left( 1-\mu_{A} \right)^{\left( y_{2}-y_{m} \right)}\left( \mathbb{I}\left[ \left( t_{2}-T_{J}-T_{A} \right)\leq y_{2}\leq\left( t_{2}-T_{J} \right) \right]R_{c}\left( t_{2}, y_{2} \right) \right)$.(S5)

Given an adult male collected on day *t_1_*, the probability that an adult mosquito collected on day *t_2_* is his offspring is expressed as

$P_{FOA}\left( t_{2}|t_{1} \right)=\frac{E_{FOA}\left( t_{2}|t_{1} \right)}{E_{A}}.$	(S6)

**Full-sibling adult (FSAA) kinship probability.** To estimate $E_{FSAA}\left( t_{2}|t_{1} \right)$, the expected number of surviving adult full-siblings on day *t_2_* of a single adult mosquito collected on day *t_1_*, we sum over all possible mating and oviposition events during the lifespan of the mother, from *y_m_* = *y_1_* - (*T_A_ - 1*) to *y_m_* = *y_1_*. As a female mosquito mates only a single time during her lifespan it is sufficient to address the father’s contribution by summing over all possible mating events. The same considerations apply with respect to the adult mosquito collected on day *t_2_* (offspring 2). Each oviposition event *R_C_* is multiplied by the survival probability *p_A_* of the mosquitoes at the corresponding event, yielding

$E_{FSAA}\left( t_{2}|t_{1} \right)=\sum_{y_{1}=t_{1}-T_{J}-\left( T_{A}-1 \right)}^{t_{1}-T_{J}} p_{A}\left( t_{1}-{y_{1}-T_{J}} \right)\sum_{y_{m}=y_{1}-\left( T_{A}-1 \right)}^{y_{1}} p_{A}\left( y_{1}-y_{m} \right)E_{FSAA}\left( t_{2}|t_{1}, y_{1},y_{m} \right).$ (S7)

$E_{FSAA}\left( t_{2}|t_{1},y_{1},y_{m} \right)= \sum_{y_{2}=y_{m}}^{y_{m}+\left( T_{A}-1 \right)} \left( 1-\mu_{A} \right)^{\left( y_{2}-y_{m} \right)}\left( \mathbb{I}\left[ \left( t_{2}-T_{J}-T_{A} \right) \leq y_{2}\leq\left( t_{2}-T_{J} \right) \right]R_{c}\left( t_{2}, y_{2} \right) \right).$(S8)

Given an adult sampled on day *t_1_*, the probability that an adult sampled on day *t_2_* is a full-sibling is expressed as

$P_{FSAA}\left( t_{2}|t_{1} \right)=\frac{E_{FSAA}\left( t_{2}|t_{1} \right)}{E_{A}}$.	(S9)

**Supplemental Text S2. Type-specific pseudo-likelihoods.**

**Adult-adult mother-offspring pseudo-likelihood**. The probability that an adult sampled on day *t*_2_ has a mother sampled on day *t*_1_ can be estimated using the previously defined kinship probability, *P_MOA_*(*t*_2_*|t*_1_), and number of adult females sampled on day *t*_1_, *n_F_*(*t*_1_), to obtain

$p_{MOA}\left( t_{2}|t_{1} \right) = 1-\left( 1-P_{MOA}\left( t_{2}|t_{1} \right) \right)^{n_{F}\left( t_{1} \right)}.$	(S10)

Because the clutch failure rate $\mu_{S}$ leads to an inflated probability of zero surviving offspring, we assume a zero-inflated binomial distribution, and define *k_MOA_* as the observed number of mother-offspring pairs among *n_A_* adults collected on day *t*_2_. The pseudo-likelihood that $k_{MOA}\left( t_{2}|t_{1} \right)$ of the *n_A_*(*t*_2_) adults sampled on day *t_2_* have a mother among the adult females sampled on day *t*_1_ is then

$L\left( k_{MOA}\left( t_{2}|t_{1} \right) \right)=ZeroInflatedBinomial\left( k_{MOA}\left( t_{2}|t_{1} \right):n_{A}\left( t_{2} \right), p_{MOA}\left( t_{2}|t_{1} \right), \mu_{S} \right).$	(S11)

The full log-pseudo-likelihood for mother-adult offspring pairs was estimated by summing over all sampling days *t_1_* and *t_2_*, or

$\Lambda_{MOA}= \sum_{t_{1}} \sum_{t_{2}} log\left( L\left( k_{MOA}\left( t_{2}|t_{1} \right) \right) \right).$	(S12)

In cases where the mother-offspring pair were collected on the same day (*t_1_* = *t_2_*), and similarly for father-offspring pairs treated below, the number of total collected adults (*n_A_*) was reduced by one to account for the fact that an adult cannot be its own parent.

**Adult-adult father-offspring pseudo-likelihood**. The probability that an adult sampled on day *t_2_* has a father sampled on day *t_1_*, can be estimated using the previously defined kinship probability, *P_FOA_*(*t*_2_*|t*_1_), and number of adult males sampled on day t1, *n_M_*(*t*_1_), to obtain

$p_{FOA}\left( t_{2}|t_{1} \right) = 1-\left( 1-P_{FOA}\left( t_{2}|t_{1} \right) \right)^{n_{M}\left( t_{1} \right)}.$	(S13)

As before, we assume a zero-inflated binomial distribution owing to a clutch failure rate $\mu_{S}$, with *k_FOA_* denoting the number of father-offspring pairs observed among *n_A_* collected adults on day *t_2_*. The pseudo-likelihood that $k_{FOA}\left( t_{2}|t_{1} \right)$ of the *n_A_(t_2_)* adults sampled on day *t_2_* have a father amongst the adult males sampled on day *t_1_* follows from the binomial distribution according to

$L\left( k_{FOA}\left( t_{2}|t_{1} \right) \right)=ZeroInflatedBinomial\left( k_{FOA}\left( t_{2}|t_{1} \right):n_{A}\left( t_{2} \right), p_{FOA}\left( t_{2}|t_{1} \right), \mu_{S} \right).$	(S14)

The full log pseudo-likelihood for father-offspring pairs was calculated by summing over all sampling days *t_1_* and *t_2_* to obtain

$\Lambda_{FOA}= \sum_{t_{1}} \sum_{t_{2}} log\left( L\left( k_{FOA}\left( t_{1}|t_{2} \right) \right) \right).$	(S15)

**Adult-adult full-sibling pseudo-likelihood.** The full log pseudo-likelihood for full-sibling pairs was estimated using the binomial distribution without zero-inflation, because the impact of clutch failure rate was already accounted for during estimation of *P_FSAA_* (Equation S9) when using $R_{c}\left( t_{2}, y_{2}, y_{1} \right)$ (Equation 1, main text). The binomial distribution was defined using the predicted number of full-sibling pairs on day *t_2_* (*k_FSAA_*), for the i-th adult mosquito sampled *s_i_*, in *n_A_* number of total collected adults on day *t_2_*, where *P_FSAA_* represents the probability to observe full-sibling pairs according to

$L\left( k_{FSAA}\left( s_{i},t_{2} \right) \right)=Binomial\left( x=k_{FSAA}\left( s_{i},t_{2} \right) {: n}_{A}\left( t_{2} \right), P_{FSAA}\left( t_{1}\left( i \right),t_{2} \right) \right).$	(S16)

The full log pseudo-likelihood for full-sibling pairs was calculated by summing over all sampled adult mosquitoes s_i_ avoiding double counting of full-sibling pairs

$\Lambda_{FSAA}= \sum_{s_{i}} \sum_{t_{2}} log\left( L\left( k_{FSAA}\left( s_{i},t_{2} \right) \right) \right).$	(S17)

**Supplemental Text S3. Prior distributions for estimated parameters.**

Prior distributions were specified for all estimated parameters. For the full model, this includes adult female census population size, juvenile mortality rate, adult mortality rate, and breeding site failure probability. Values of other model parameters were fixed at assumed values of β = 40 eggs per day, *T_J_* = 15 days, and *T_A_* = 25 days.

The prior distribution for adult female census population size on Jaana Island was established using a combination of published estimates of *An. gambiae* effective population size on the Ssese Islands (Kayondo et al. 2005; Wiltshire et al. 2018; Bergey et al. 2020) and estimates of the ratio of effective population size (*N_e_*) and census population size (*N*) (Palstra and Fraser 2012). Random samples were drawn from the published values of *N_e_* and the *N_e_/N* ratio, which we then used to calculate values of *N* and summarized with a log-normal distribution with parameter values obtained using maximum likelihood. The resulting prior distribution for *N_F_* had a mean of 4,317 individuals and a 95% uncertainty interval of 252 to 77,971 individuals.

The prior distributions for juvenile and adult mortality rates were specified by fitting a normal distribution to logit-transformed central estimates of daily survival probabilities from a number of published mosquito survival experiments (Service 1965, 1971, 1973; Costantini et al. 1996; Bayoh and Lindsay 2004; Munga et al. 2007; Matthews et al. 2020). This resulted in an average mortality rate of 0.171 (95% uncertainty interval: 0.02 - 0.39) for juveniles and 0.109 (95% uncertainty interval: 0.03 - 0.28) for adults.

The prior distribution of the breeding site failure probability was estimated by first obtaining 10^4^ random samples from the prior distributions of the juvenile and adult mortality rates. Then, assuming a constant population size (which is a simplistic but reasonable assumption for the long-term dynamics of a natural population), the demographic parameters are related to one another as

$1= \beta\left( 1-\mu_{J} \right)^{T_{J}}\left( 1-\mu_{S} \right)^{T_{J}}\sum_{i}^{T_{A}} \left( 1-\mu_{A} \right)^{i}$.	(S18)

Leveraging this relationship and the specified values of all the other parameters, we calculated corresponding values of μ_S_. This resulted in 10^4^ values of μ_S_ that were tied to 10^4^ values of μ_J_ and μ_A_, which were then input into Bayesian Tools for MCMC sampling using the createPriorDensity function.

For the model in which $\mu_{S}$ was fixed at zero, we used a similar, but modified, approach to specifying prior distributions. In this case, the prior for $\mu_{A}$ was the same as in the full model. For $\mu_{J}$, we applied a similar procedure as was used to specify the prior for $\mu_{S}$ in the full model. Specifically, we drew 10^4^ random samples from the prior distribution of $\mu_{A}$. Then, we applied Equation S18 with $\mu_{S}$ set to zero to solve for $\mu_{J}$. Repeating this for each of the 10^4^ random samples of $\mu_{A}$ yielded 10^4^ corresponding values of $\mu_{J}$ that defined its prior distribution.

**Supplemental Text S4. *Anopheles gambiae* life history characteristics**

*Anopheles gambiae* development is holometabolous, with three aquatic juvenile (immature) stages (egg, larva, pupa) followed by the terrestrial adult stage. Although the rate of development is temperature-dependent, eggs normally take 1-2 days to hatch, the larval stage lasts 6-9 days, and the pupal stage 1-2 days. A minimum duration from egg to adult may be 8 days, but because an additional ~4 days are typically required for the production of the first batch of eggs, a minimum generation time is 10-11 days, often 1-2 days more (Gillies and De Meillon 1968). Adult females can live up to a month but the majority live 2 weeks or less in nature. Based on mark-recapture experiments involving other anopheline species, it was estimated that the average lifespan of wild males was 5-10 days (Howell and Knols 2009 and references therein). Adult emergence occurs in the early evening, and is followed by a teneral resting period of ~24 hours during which the cuticle hardens (Clements 1999). During this same period, male structures essential for sexual activity mature: the male terminalia rotates 180° to enable proper orientation for mating, and the antennal fibrillae acquire full functionality (Howell and Knols 2009). Following 1-2 days of sexual maturation post-emergence, males form swarms around dusk, and females fly into the swarms to mate. Once mated, a female typically is no longer receptive to further mating during her lifetime, but she is iteroparous. Sperm stored in the spermatheca fertilize large clutches of eggs (between 50 and 300) that require a blood meal to develop. After obtaining a full blood meal, the female rests 2-3 days while the blood is digested and eggs are developed, followed by oviposition. The cycle of blood feeding and oviposition repeats until the female dies, potentially having laid 800-1000 eggs during her lifetime. Although males may mate multiple times, many or most do not have the opportunity to do so, and it is estimated that the typical male mates 0-3 times (Howell and Knols 2009).

**Supplemental Text S5. Individual-based simulation model description.**

The simulations assumed equal sex ratio at oviposition, age-invariant fecundity and adult survival probability, constant adult population abundance, and no immigration or emigration, to match the assumptions used in the CKMR model. At the same time, the individual-based model differed from the CKMR model in the following ways to more realistically represent *An. gambiae* biology on Jaana Island. First, in the individual-based model, the initial oviposition of the female occurs four days after emergence (one day mating, one day blood feeding, and two days resting), whereas oviposition in the CKMR model begins right after emergence. This difference led to an overall reduction in oviposition over the lifespan of a female mosquito. Second, in the individual-based model, the female mosquito oviposited every three days and the number of eggs per oviposition β was drawn from a Poisson distribution with mean 120. In the CKMR model, the female mosquito oviposited 40 eggs daily, a feature of the Sharma et al. (2022) formulation that was made for computational ease. Third, in the individual-based simulation, emergent females mated a single time randomly with adult male mosquitoes.

To the extent possible given differences in certain assumptions between the Sharma et al. CKMR model and the individual-based SLiM simulation model, we configured our simulations to match the conditions of the PO+FS CKMR model fitted to observed data from Jaana Island (Table 2). To simulate adult dynamics, we took a Bernoulli random draw of each individual adult’s mortality with probability 𝜇_A_ on each day. Recruitment of new adults occurred automatically for any individual juveniles who survived for *T_j_* days. To simulate juvenile dynamics, additional modifications were required to ensure that the behavior of stochastic simulations was consistent with expectations of the CKMR model. The deterministic formulation of the CKMR likelihood assumes that mortality and recruitment occur at fixed rates, which would result in significant drift of the population size over time if implemented this way in a stochastic model. To guard against that, it was necessary to introduce a mechanism for density dependence into the individual-based SLiM simulation model. To do this, we set a carrying capacity, *C*, for juvenile mosquitoes. Whenever the total number of simulated juveniles on day *t*, *N_J,t_*, was below *C*, individuals survived with probability 1– *N_J,t_* / *C*. Consistent with our interest in exploring clutch-level mortality events through breeding site failure, this mortality outcome was applied to all individuals within a clutch. The value of *C* was obtained through calibration to achieve a desired adult female population size, *N_F_*. In addition to *C*, we initialized the simulation by adding *N_α_* emerging adult mosquitoes daily during the first 19 days of the simulation, at which time the first adults arising from simulated ovipositions emerged (a time span determined by one day for mating, one day for blood feeding, two days for resting, and 15 days of juvenile maturation time *T_J_*). The number of emerging adult mosquitoes *N_α_* was calculated by dividing the adult female population size *N_F_* by the summed daily survival, and multiplied by two assuming equal sex ratio at emergence, resulting in

$N_{\alpha}=\frac{N_{A}}{\sum_{i=1}^{T_{A}} \left( 1-\mu_{A} \right)^{i}}$.	(S19)

We confirmed through simulation that the chosen values of *C* and *N_α_* resulted in the desired number of simulated adult females, *N_F_*.

After a burn-in period of 100 days iteration for each simulation scenario, we mimicked the actual 20-day sampling of adult mosquitoes on Jaana Island over a further 20-day iteration before simulations were ended at 121 days. Thus, the same number of blood fed females, females without blood meals, and male mosquitoes were randomly and lethally sampled during the population simulation, on the corresponding collection days recorded in the actual field study (Table S5). The resulting simulated field collection matched in size and composition on each of 20 collection days the field collection studied with the CKMR models, but in the simulated field collection the true pairwise kinship relationships and the true abundance of the simulated adult population being sampled was known.

**Supplemental Text S6. Estimating the migration rate.**

Our rough estimate of the migration rate was derived as follows. In a single closed constant-size population with discrete generations, the probability that two individuals chosen at random are full sibs is $\sigma^{2}/N_{F}$, where $\sigma^{2}$ is the variance of female reproductive success and $N_{F}$ is the number of females. If $n$ individuals are sampled, then there are approximately $n^{2}/2$ pairs, and thus the expected number of full sib pairs will be the product of the two. When there are two demes, if we assume (1) a fraction $m$ of individuals in each deme are immigrants from the other population; (2) individuals can only migrate before they reproduce (so full sibs are always born in the same deme); and (3) sampling occurs after any movement, then the expected number of full sib pairs in the first deme will be

$FS_{1}=\frac{\sigma^{2}n_{1}^{2}}{2}[\frac{\left( 1-m \right)^{2}}{N_{1}}+\frac{m^{2}}{N_{2}}]$, and analogously for the second deme, while the expected number of full sib pairs between demes will be ${{FS}_{12}=\sigma}^{2}n_{1}n_{2}m(1-m)(\frac{1}{N_{1}}+\frac{1}{N_{2}})$. These equations can be rearranged to give $2m\left( 1-m \right)=\frac{{FS}_{12} r}{{FS}_{1}+{FS}_{12}r+{FS}_{2}r^{2}}$, where $r={n_{1}}/{n_{2}}$ is the ratio of the two sample sizes. If we replace these expectations with our observed values ${(FS}_{1}=36; {FS}_{12}=6; {FS}_{2}=18; r=613/101)$, and solve for $m$, we obtain $m=0.025.$ Similar rearrangements allow us to calculate the ratio of the population size at LWZ to that at KKU/NKD as ${N_{1}}/{N_{2}=19.4}$. These estimates (which do not come with confidence limits) should be taken as indicative only; future work could expand the more formal MCMC approach we applied to LWZ to consider 2 demes.

**Supplemental Text S7. Estimating the variance of female and male reproductive success and the effective population size (N_e_).**

Here we derive a number of expressions for the mean and variance of various components of reproductive success and for N_e_.

*Number of surviving clutches for a female.*

The CKMR model assumes that females have a constant daily mortality rate $m_{A}$, and that each day she is alive she lays a clutch, with egg laying occurring before death, so the number $C$ of clutches laid by a female before she dies will be

$$C\sim geometric\left( \mu_{A} \right)$$

[S20]

with pmf

$$f_{C}(c)=\left( 1-\mu_{A} \right)^{c-1}\mu_{A}$$

Each clutch has a daily probability of failing of $\mu_{S}$, and eggs take 15 days to mature into adults, so the overall probability of a clutch failing is:

$$\mu_{C}=1-\left( 1-\mu_{S} \right)^{15}$$

[S21]

If $S_{F}$ is the number of surviving clutches from a female, then conditionally, $S_{F}|C$ follows a binomial distribution:

$$S_{F}|C\sim binomial(C, 1-\mu_{C})$$

The marginal distribution of $S_{F}$ is

$$f_{S_{F}}\left( s \right)=\frac{\mu_{A}\left( 1-\mu_{A} \right)^{s-1}\left( 1-\mu_{C} \right)^{s}}{\left[ 1-\left( 1-\mu_{A} \right)\mu_{C} \right]^{s+1}}$$

for $S_{F}\geq1$, and for $S_{F}=0$ we have

$$f_{S_{F}}\left( s \right)=\frac{\mu_{A}\mu_{C}}{1-\left( 1-\mu_{A} \right)\mu_{C}}$$

[S22]

We can calculate the mean and variance of the number of surviving clutches from [S20] or via the total law of expectation / variance:

$$E\left[ S_{F} \right]=\frac{1-\mu_{C}}{\mu_{A}}$$

$$Var\left[ S_{F} \right]=\frac{\mu_{C}\left( 1-\mu_{C} \right)}{\mu_{A}}+\frac{\left( 1-\mu_{C} \right)^{2}\left( 1-\mu_{A} \right)}{\mu_{A}^{2}}$$

[S23]

*Number of offspring from a female*

If each surviving clutch produces surviving offspring according to a Poisson distribution with rate lambda, then the total number of offspring $T_{F}$ from a female will also be distributed as Poisson (sum of independent Poisson’s):

$$T_{F}|S_{F}\sim Poisson(\lambda S_{F})$$

The model assumes constant population size, so females must on average produce 2 surviving offspring, and therefore the number of surviving offspring per surviving clutch will be

$$\lambda=\frac{2}{E\left[ S_{F} \right]}=\frac{2\mu_{A}}{1-\mu_{C}}$$

It is easy to verify that $E\left[ T_{F} \right]=2$:

$$E\left[ T_{F} \right]=E\left[ E\left( T_{F} | S_{F} \right) \right]=E\left[ \lambda S_{F} \right]=\lambda E\left[ S_{F} \right]=2$$

The variance of the number of surviving offspring from a female will then be:

$$Var\left[ T_{F} \right]=E\left[ \lambda S_{F} \right]+Var\left[ \lambda S_{F} \right]$$

$$=2+\lambda^{2}Var\left[ S_{F} \right]$$

$$=2+4\left[ \frac{\mu_{A}\mu_{C}}{1-\mu_{C}}+1-\mu_{A} \right]$$

[S24]

*Number of mates of a male*

The model assumes that adult males have the same daily mortality rate as females, and thus their adult lifespan is also distributed as $C$ in [20]. Females mate only once in their life, and since there is an equal sex ratio, males also on average mate once in their life but with variation. Therefore, the daily probability a male mates will be equal to the reciprocal of the mean lifespan. If we let the number of females mated by a male be $K$, then

$$K|C\sim binomial(C, \mu_{A})$$

Again, we can confirm there is there is one mating event on average:

$$E\left[ K \right]=E\left[ E\left( K | C \right) \right]=E\left[ N\mu_{A} \right]=1$$

The variance will then be:

$$Var\left[ K \right]=2\left( 1-\mu_{A} \right)$$

[S25]

*Number of surviving clutches sired by a male*

As a male can mate $K$ times then the total number of surviving clutches $S_{M}$ is the sum of $K$ independent $S_{F}$’s:

$$S_{M}|K\sim\sum^{K} S_{F_{i}}$$

Of course when $K=0$ then $S_{M}=0$. Its mean and variance are:

$$E\left[ S_{M} \right]=E\left[ K \right]E\left[ S_{F} \right]$$

$$Var\left[ S_{M} \right]=E\left[ K \right]Var\left[ S_{F} \right]+E\left[ S_{F} \right]^{2}Var[K]$$

Since males mate on average once ($E\left[ K \right]=1$), $E\left[ S_{M} \right]=E[S_{F}]$. For the variance,

$$Var\left[ S_{M} \right]=\frac{\mu_{C}(1-\mu_{C})}{\mu_{A}}+\frac{3\left( 1-\mu_{C} \right)^{2}(1-\mu_{A})}{\mu_{A}^{2}}$$

[S26]

*Number of offspring of a male*

Similarly, the total number of offspring $T_{M}$ given by a male is the sum of $K$ independent $T_{F}$’s:

$$T_{M}|K\sim\sum^{K} T_{F_{i}}$$

with mean and variance:

$$E\left[ T_{M} \right]=E\left[ K \right]E[T_{F}]$$

$$Var\left[ T_{M} \right]=E\left[ K \right]Var\left[ T_{F} \right]+E\left[ T_{F} \right]^{2}Var[K]$$

As before, we assume $E[K]=1$ and females have on average 2 offspring ($E[T_{F}]=2)$, this becomes

$$Var\left[ T_{M} \right]=Var\left[ T_{F} \right]+4Var[K]$$

In other words, mating variance brings an additional $4Var[K]$ to the variance of reproductive success. Using [24] and [25], we then have:

$$Var\left[ T_{M} \right]=2+4\left[ \frac{\mu_{A}\mu_{C}}{1-\mu_{C}} \right]+12\left( 1-\mu_{A} \right)$$

[S27]

*Effective population size*

From Crow and Denniston (1988) Eqn 33 we have a formula to calculate effective population size $N_{eV}$:

$$\frac{1}{N_{eV}}\approx\frac{1}{4N_{M}}\left[ f+m\frac{Var\left( T_{M} \right)}{E\left( T_{M} \right)} \right]+\frac{1}{4N_{F}}[m+f\frac{Var\left( T_{F} \right)}{E\left( T_{F} \right)}]$$

where $N_{M}$ and $N_{F}$ are the number of males and female adults, and $m+f=1$ are the proportions of male and female progeny. The subscript $V$ indicates it is a variance effective population size, which relates the magnitude of genetic drift of the focal population to an idealised one. Further, if equal sex-ratio and constant population size are assumed, such that $N_{M}=N_{F}=N$, $m=f=0.5$, and $E[T_{M}]=E[T_{F}]=2$, we have

$$\frac{1}{N_{eV}}\approx\frac{1}{4N}[1+\frac{Var\left( T_{F} \right)}{2}+Var\left( K \right)]$$

Further simplification gives

$$\frac{1}{N_{eV}}\approx\frac{1}{2N}\left[ 1+\frac{\mu_{A}\mu_{C}}{1-\mu_{C}}+2\left( 1-\mu_{A} \right) \right]$$

[S28]

The various formulae for means, variances, and parameter estimates are summarized in the table below:

| **Variable** | **Symbol** | **Theoretical expectation and estimate (from LWZ data)** | | **Theoretical variance and estimate (from LWZ data)** | |
| --- | --- | --- | --- | --- | --- |
| Clutches produced per female | $C$ | $E\left[ C \right]=\frac{1}{\mu_{A}}$ | 5.70  [3.84, 10.39] | $Var\left[ C \right]=\frac{1-\mu_{A}}{\mu_{A}^{2}}$ | 26.81  [10.89, 97.47] |
| Surviving offspring per clutch | $\lambda$ | $\lambda=\frac{2\mu_{A}}{1-\mu_{C}}$ | 15.21  [3.97, 84.01] | $\lambda=\frac{2\mu_{A}}{1-\mu_{C}}$  [Assumed to be Poisson] | - |
| Surviving clutches per female | $S_{F}$ | $E\left[ S_{F} \right]=\frac{1-\mu_{C}}{\mu_{A}}$ | 0.131  [0.024, 0.503] | $Var\left[ S_{F} \right]=\frac{\mu_{C}\left( 1-\mu_{C} \right)}{\mu_{A}}+\frac{\left( 1-\mu_{C} \right)^{2}\left( 1-\mu_{A} \right)}{\mu_{A}^{2}}$ | 0.14  [0.024, 0.658] |
| Surviving offspring per female | $T_{F}$ | $E\left[ T_{F} \right]=2$ [model assumption] | - | $Var\left[ T_{F} \right]=2+4\left[ \frac{\mu_{A}\mu_{C}}{1-\mu_{C}}+1-\mu_{A} \right]$ | 35.0  [12.5 – 172.6] |
| Males mated per female | - | 1 [model assumption] | - | - | - |
| Females mated per male | $K$ | $E[K]=1$ [model assumption] | - | $Var\left[ K \right]=2\left( 1-\mu_{A} \right)$ | 1.65  [1.48, 1.81] |
| Surviving clutches sired per male | $S_{M}$ | $E\left[ S_{M} \right]=\frac{1-\mu_{C}}{\mu_{A}}$ | Same as $E[S_{F}]$ | $Var\left[ S_{M} \right]=\frac{\mu_{C}\left( 1-\mu_{C} \right)}{\mu_{A}}+\frac{{3\left( 1-\mu_{C} \right)}^{2}\left( 1-\mu_{A} \right)}{\mu_{A}^{2}}$ | 0.170  [0.025, 1.065] |
| Surviving offspring per male | $T_{M}$ | $E\left[ T_{M} \right]=2$ [model assumption] | - | $Var\left[ T_{M} \right]=2+4\left[ \frac{\mu_{A}\mu_{C}}{1-\mu_{C}} \right]+12\left( 1-\mu_{A} \right)$ | 41.6  [19.0 – 179.3] |

The parameter estimates are based on the posterior distributions of $\mu_{A}$ and $\mu_{S}$ from which the medians are reported. The 95% C.I.s are calculated from the 2.5- and 97.5-percentile of the posteriors of the parameters of interest.
